# The restriction factor *pastrel* is associated with host vigor, viral titer, and variation in disease tolerance during Drosophila C Virus infection

**DOI:** 10.1101/2022.06.09.495537

**Authors:** Megan A.M. Kutzer, Vanika Gupta, Kyriaki Neophytou, Vincent Doublet, Katy M. Monteith, Pedro F. Vale

**Affiliations:** Institute of Evolutionary Biology, School of Biological Sciences, University of Edinburgh, UK; Institute of Immunology and Infection Research, School of Biological Sciences, University of Edinburgh, UK; Institute of Evolutionary Ecology and Conservation Genomics, University of Ulm, Germany

## Abstract

Genetic variation for both resistance and disease tolerance has been described in a range of species infected with bacterial, viral and fungal pathogens. In *Drosophila melanogaster*, genetic variation in mortality following systemic Drosophila C Virus (DCV) infection has been shown to be driven by large effect polymorphisms in the viral restriction factor *pastrel (pst)*. However, it is unclear if *pst* impacts variation in DCV titres (i.e. resistance), or if it also contributes to disease tolerance. We investigated systemic infection across a range of DCV challenge doses spanning nine orders of magnitude, in males and females of ten *Drosophila* Genetic Reference Panel (DGRP) lines carrying either a susceptible (S) or resistant (R) *pst* allele. Our results uncover among-line variation in fly survival, viral titers, and disease tolerance measured both as the ability to maintain survival (mortality tolerance) and reproduction (fecundity tolerance). We confirm the role of *pst* in resistance, as fly lines with the resistant (R) *pst* allele experienced lower viral titers, and we uncover novel effects of *pst* on host vigor, as flies carrying the R allele exhibited higher survival and fecundity even in the absence of infection. Finally, we found significant variation in the expression of the JAK-STAT ligand *upd3* and the epigenetic regulator of JAK-STAT *G9a*. While *G9a* has been previously shown to mediate tolerance of DCV infection, we found no correlation between the expression of either *upd3* or *G9a* on fly tolerance or resistance. Our work highlights the importance of both resistance and tolerance in viral defence.

## Introduction

Why do some hosts succumb to infection while others survive? Host heterogeneity in infection outcomes can be attributed in part to two distinct but complimentary sets of mechanisms, which together act to maintain host health: mechanisms that limit pathogen growth and mechanisms that prevent, reduce or repair tissue damage caused during infection but without directly affecting pathogen load. The relative balance between these mechanisms may result in phenotypically distinct outcomes. We tend to associate a strong capacity to clear infection with a ‘resistance’ phenotype, while hosts with very efficient damage limitation mechanisms may appear to be relatively healthy even if their ability to clear is not pronounced and pathogen loads remain high - generally described as a ‘disease tolerance’ phenotype [1–7].

Beyond differences in their underlying mechanisms, resistance and tolerance can have profoundly different epidemiological and evolutionary outcomes [8–11]. If disease tolerance improves host survival, the infectious period is prolonged, thus increasing pathogen transmission and infection prevalence. In this case, hosts with an allele that confers mortality tolerance (high survival relative to their pathogen load) have a fitness advantage, so the tolerance allele spreads throughout the host population, leading to the eventual fixation of tolerance in the population [12]. However, this prediction contrasts with many studies that find evidence for genetic variation in disease tolerance within a population [13–17]. On possible explanation for this divergence between predicted and observed levels of genetic variation is that disease tolerance may incur fitness costs that are not captured in models of tolerance evolution. A related point is that many evolutionary models make specific assumptions about a trade-off between resistance and tolerance [12,18]. While such a trade-off may exist in some systems [13,16], it is by no means universal [19–21]. Further, if disease tolerance acts only to maintain or improve host fecundity, it should be neutral with respect to pathogen prevalence because host lifespan is unaffected, thus the pathogen’s transmission period is neither prolonged nor shortened [12]. Therefore, theoretical predictions suggest that we might expect to observe heterogeneity for fecundity tolerance but not mortality tolerance in natural populations [12].

Here, we tested how two intrinsic sources of variation – genetic background and sex – interact to contribute to host heterogeneity in disease defence measured as resistance and tolerance. We focused on the interaction between the fruit fly *Drosophila melanogaster* and Drosophila C Virus (DCV), a horizontally transmitted, positive sense RNA virus, that naturally infects multiple *Drosophila* species [22–24]. Systemic infection with high doses of DCV leads to infection of the smooth muscles around the crop, which causes pathology and results in intestinal obstruction, reduced metabolic rate, and reduced locomotor activity [25–28]. The majority of genetic variance in host mortality during DCV infection is controlled by large effect polymorphisms in and around the *pastrel* (*pst*) gene, a viral restriction factor [22,29]. The protective effect of *pst* was confirmed by loss-of-function mutants and an overexpression study [29]. However, it is unclear if variation in the protective effects of *pst* act by increasing the fly’s ability to clear the viral infection, or to tolerate its pathological effects. Further, DCV infection is associated with increased fecundity as well as accelerated developmental time in larvae at both lethal and sublethal doses [26]. Since *D. melanogaster* may tolerate infections by increasing their reproductive output and/ or improving survival outcomes, we used lines that varied in their susceptibility to DCV infection [30] in order to capture the entire range of genetic variation in resistance and tolerance available across the DGRP panel.

We used males and females flies from ten DGRP lines [31] carrying either a resistant (R) or susceptible (S) *pst* allele. We systemically challenged male and female flies with five doses of DCV. We measured host lifespan and viral titre in both sexes, as well as cumulative fecundity and reproductive rate in females. By doing so, we were able to characterize natural variation in resistance, mortality tolerance, and fecundity tolerance to DCV. Tolerance is frequently measured as a reaction norm, where host fitness is regressed against parasite load assayed at a fixed dose [1,4]. Instead of relying on host heterogeneity at a single dose, we regressed host lifespan and cumulative fecundity against five viral doses spanning nine orders of magnitude to examine variation in mortality and fecundity tolerance (see also [32,33]. This allowed us to assess how each fly genotype and sex contribute to host defence across a broad range of infection intensities.

In addition to characterizing variation in resistance to and tolerance of DCV infection, we also aimed to link this variation with potential mechanisms, particularly for disease tolerance, where knowledge of the underlying mechanisms has lagged behind the description of their phenotypic effects. As disease tolerance relates to a reduction of pathology independently of pathogen clearance, tolerance mechanisms described to date have included those that prevent, limit or repair tissue damage [3,34–38]. Inflammation is one common cause of such damage during infection. Pro-inflammatory cytokines tend to be associated with decreased tolerance to infection – for example, a tolerant house finch population (*Haemorhous mexicanus*) infected with a bacterial pathogen, *Mycoplasma gallisepticum*, exhibited lower cytokine expression compared with a less tolerant population [35]; mice receiving the anti-inflammatory drug Ibuprofen showed improved increased tolerance during *Mycobacterium tuberculosis* infection [39]; and lower levels of circulating pro-inflammatory cytokines are associated with tolerance of malaria after re-exposure to the parasite [40]. Negative regulation of immune responses that minimize inflammation would therefore appear to be prime candidates for mechanisms that promote disease tolerance [34,40–42]. This is supported by previous work showing that the epigenetic modifier, *G9a*, which regulates JAK-STAT signalling to prevent hyperactivation of the immune response, increases tolerance to RNA virus infection by limiting immunopathology [43,44]. We therefore also investigated if variation in resistance or tolerance in the tested lines were associated with the expression of either *G9a* or of *upd3*, a JAK-STAT pathway target gene that encodes a cytokine-like protein [45].

## Methods

### D. melanogaster culture conditions and experimental lines

To assess genetic variation in resistance and tolerance to *Drosophila* C virus (DCV), we chose ten lines from the *Drosophila* Genetic Reference Panel (DGRP) [31] spanning the range of variation in fly survival within the DGRP when infected systemically with DCV [22]. Because the transcription factor *pastrel* (*pst)* is known to affect survival to DCV infection, we specifically selected five susceptible (S) lines (RAL-138, RAL-373, RAL-380, RAL-765, RAL-818) and five resistant (R) lines (RAL-59, RAL-75, RAL-379, RAL-502, RAL-738). All lines were previously cleared of *Wolbachia* infection, as it known to confer protection against DCV. All fly stocks in the lab, including the DGRP panel, are routinely checked for several viral pathogens using PCR; no viral contamination has ever been detected [46–48]. All lines were maintained on standard cornmeal medium (cite) at 25°C on a 12h: 12h light: dark cycle.

### Virus preparation

DCV was grown in a *Drosophila* S2 cell culture as described previously [27]. The homogenized culture was passed through a sucrose cushion, ultracentrifuged and re-suspended in 10mM Tris-HCl (pH 7.3). The suspended virus was stored at -80°C in 10 µl aliquots. Virus titres were measured using quantitative Real Time PCR as described previously [44]. Briefly, total RNA was extracted using TRI reagent (Ambion) and then reverse transcribed using M-MLV Reverse Transcriptase (Promega) and random hexamers. The manufacturer’s protocol was followed to synthesize cDNA. Ten-fold serial dilutions of this cDNA was done up to 10^−10^ dilution. The number of DCV copies in these samples was quantified using DCV specific primers (DCV_Forward: 5′ AATAAATCATAAGCCACTGTGATTGATACAACAGAC 3′, DCV_Reverse: 5′ AATAAATCATAAGAAGCACGATACTTCTTCCAAACC 3′) and Fast SYBR green (Applied Biosystems) based qRT-PCR (Applied Biosystems StepOne Plus). The dilution at which no copies were detected was set as zero reference. The viral quantity was back calculated from this point and viral copies in the stock were estimated to be 10^9^ DCV infective units (IU) ml^-1^.

### Infections

All experimental flies were reared under constant density of between 80-100 eggs per vial for at least two generations. We infected 3 – 5 day old adult male and female flies with five concentrations of DCV inoculum 10^3^, 10^5^, 10^6^, 10^8^, and 10^9^ DCV IU ml^-1^. All the viral inoculums were obtained by diluting the same viral stock solution in sterile 10mM Tris-HCl. Flies were infected systemically by intra-thoracic pricking using a needle (Minutein pin, 0.14mm) dipped in the viral suspension. A control group were pricked with a needle dipped in sterile 10mM Tris-HCl (pH - 7.3). In total, we infected 20 individual replicate flies for each combination of DGRP line, DCV concentration and sex, resulting in a total of 2400 flies (20 replicates x 10 DGRP lines x 6 DCV concentrations x 2 sexes). Given the large number of infections (5 replicates per Line x Dose x Sex, ∼ 600 flies per day), we blocked the experiment across four days, and collected eggs separately from each of the ten DGRP lines on each day. Each fly was housed individually in a vial after infection and flies were monitored for mortality daily. Flies were transferred to new food vials every week until day 28 post infection, while the previous vials were stored at 25°C until all progeny eclosed as adults. We quantified the cumulative fecundity of each individual fly as the total number of adult offspring produced during this 28-day period (or until death, if this happened prior to the 28^th^ day).

### Viral load

In addition to the 2400 flies exposed to DCV to monitor survival, a further five individuals for a given Line × Dose × Sex combination (600 flies in total) were infected to measure the viral load at three days post infection (3 DPI). We chose this time-point because we wanted to quantify viral load in the flies before the onset of mortality due to infection across all doses as flies in the higher DCV concentrations started dying within four days of infection. Each fly was transferred to TRI reagent at 3 DPI, and flies were frozen at -80°C until RNA extraction. We measured viral load as described above and previously in Gupta and Vale (2017). We generated the DCV standard curve by quantifying DCV titers in serially-diluted samples of DCV. This standard curve was used for absolute quantification of virus titers in the fly samples.

### Gene expression

The Jak-Stat pathway has been described previously as being involved in the response to DCV [43]. To test if measures of resistance or tolerance were correlated with the expression of Jak-Stat pathway genes, we pricked 3 – 7 day old flies with 10 mM Tris-HCl (pH 7.3) (control) or 10^7^ DCV IU ml^-1^. We used 10^7^ DCV IU ml^-1^ because it reflected the half maximal effective concentration (EC50) across the 10 tested lines and elicits an immune response in *D. melanogaster* at this dose. Following infection, the flies were housed by Line x Treatment x Sex in vials containing standard Lewis Cornmeal medium. Three days post-infection, we set up five replicates of each treatment combination containing three live flies in 1.5 mL Eppendorf tubes. We anesthetized the flies on ice, placed them in 60 μl of TRIzol reagent (Invitrogen), and stored them at -70°C for gene expression analyses.

To quantify the differences in transcription levels of *G9a* and the Jak-Stat pathway gene *upd3*, we used quantitative Reverse Transcription PCR (RT-qPCR). First, we homogenized flies submerged in TRIzol Reagent using a pestle motor. Total RNA was extracted using a Direct-zol RNA Miniprep kit (Zymo Research) in accordance with the manufacturer’s instructions and stored at -70°C. We included a DNase treatment step per the manufacturer’s recommendation, to digest genomic DNA. The isolated RNA was reverse transcribed with M-MLV reverse transcriptase (Promega) and random hexamer primers (ligation at 70°C for 5 mins, cDNA synthesis at 37°C for 1 hr), diluted 1:7 with triple-distilled water and stored at -20°C. Gene expression was quantified using Fast SYBR Green Master Mix (Applied Biosystems) and the primers detailed in Supplementary Table S1, on the Applied Biosystems StepOnePlus instrument using the following protocol: 95°C for 2 mins, followed by 40 cycles of denaturation at 95°C for 10 s and annealing and amplification at 60°C for 30 s. We normalized gene expression of the target genes with the reference gene *rp49* and reported expression as fold change relative to the control flies. We calculated fold change in gene expression as 2^-ΔΔCt[49]^.

To correct for the systematic error among qPCR plates (n = 10), we used two calibrators (male RAL-501, replicate 1, infected; male RAL-501, replicate 1, uninfected). Eight μl aliquots were stored at -20°C for later use. The calibrators’ mean Ct values were used to calculate correction factors per run, per target gene. Between-plate variation was removed prior to calculating relative gene expression, as described by [50]. Missing values for the *G9a* calibrators for one plate were determined from the correlation of *G9a* expression from all runs between calibrators’ mean Ct and Ct values of samples.

### Statistical methods

Statistical analyses were performed in R version 4.0.4 and R Studio 1.4.1106. Models 1a and 1b were analyzed with a Cox mixed effects survival model using the coxme function in the coxme package [51]. We used Gamma glms (glm function in R base stats package) to evaluate Models 2a and 2b and multiple linear regressions (lm function in the R base stats package) to evaluate Models 5a – 8b. Generalized linear mixed models (Models 3a, 3b, 4a, and 4b) were analyzed using the glmmTMB function with negative binomial error structures with a quadratic parameterization (nbinom2) for Models 3a and 3b or with a linear parameterization (nbinom1) and zero inflation for Models 4a and 4b [52]. Models 4a and 4b included lifespan as an offset term to control for its effects on cumulative fecundity. We tested for significant interactions and/or main effects using type 2 or 3 Wald χ^2^ or F tests [53] as appropriate. Experimental block was included as a random effect in Models 1a, 1b, 3a, 3b, 4a, and 4b. All models were evaluated using model selection criteria [54] and using the check_model function in the performance package if applicable. Interactions were excluded from the final models if p<0.1. Models are further described in Tables 1 – 4 and individual model parameter estimates are included Supplementary Tables S2 – S17 within Appendix S1. Correlations were assessed using Kendall’s *tau* coefficient. In Figure 3c, the y-intercept of each function was standardized at 0 to account for differences in general vigor, *e*.*g*., Råberg *et al* 2009, before integration.

**Table 1.** The effects of DCV dose, sex, and *pst* or DGRP line on lifespan and mortality tolerance. Model 1a tested survival differences between R and S *pst* alleles. Model 2a tested survival differences among DGRP lines and Models 3a and 3b tested for differences in mortality tolerance between *pst* alleles or among DGRP lines (indicated by a statistically significant interaction with dose or dose^2^). Values in bold are statistically significant.

| Response | Model # | Predictor | Df | $\chi^2$ | P-value |
| --- | --- | --- | --- | --- | --- |
| Lifespan | 1a | Dose | 1 | 948.7459 | **<0.0001** |
|  |  | Pst allele | 1 | 28.103 | **<0.0001** |
|  |  | Sex | 1 | 2.8748 | 0.0899 |
|  | 1b | Dose | 1 | 110.1135 | **<0.0001** |
|  |  | Line | 9 | 44.4518 | **<0.0001** |
|  |  | Sex | 1 | 3.2453 | 0.0716 |
| | | Dose × Line | 9 | 20.9098 | **0.0131** |
| | | Line × Sex | 9 | 29.9929 | **0.0004** |

## Results

### pastrel affects fly survival during infection and vigor in the absence of infection

First, we examined the effects of DCV dose and sex on survival across ten genotypes to determine if hosts varied in their susceptibility to viral infection. Because *pst* is known to affect fly mortality following DCV infection, we selected five S lines (138, 373, 380, 765, 818) and five R lines (59, 75, 379, 502, 738), based on previously described infected lifespans [22]. As expected, R lines tended to live longer than S lines (Figure 1a, Table 1, Model 1a, *pst* allele: p <0.0001), though this was also the case in the absence of infection (*i*.*e*., general vigor). Examining all ten lines separately, we detected genetic variation in survival and found that the ten tested lines differed in their responses to dose (Figure 1B and 1C; Table 1, Model 1b, Line x Dose: p = 0.013), while sex and genetic background affected survival independently of dose (Figure 1B and 1C; Table 1, Model 1b, Line x Sex: p = 0.0004).

**Figure 1.**
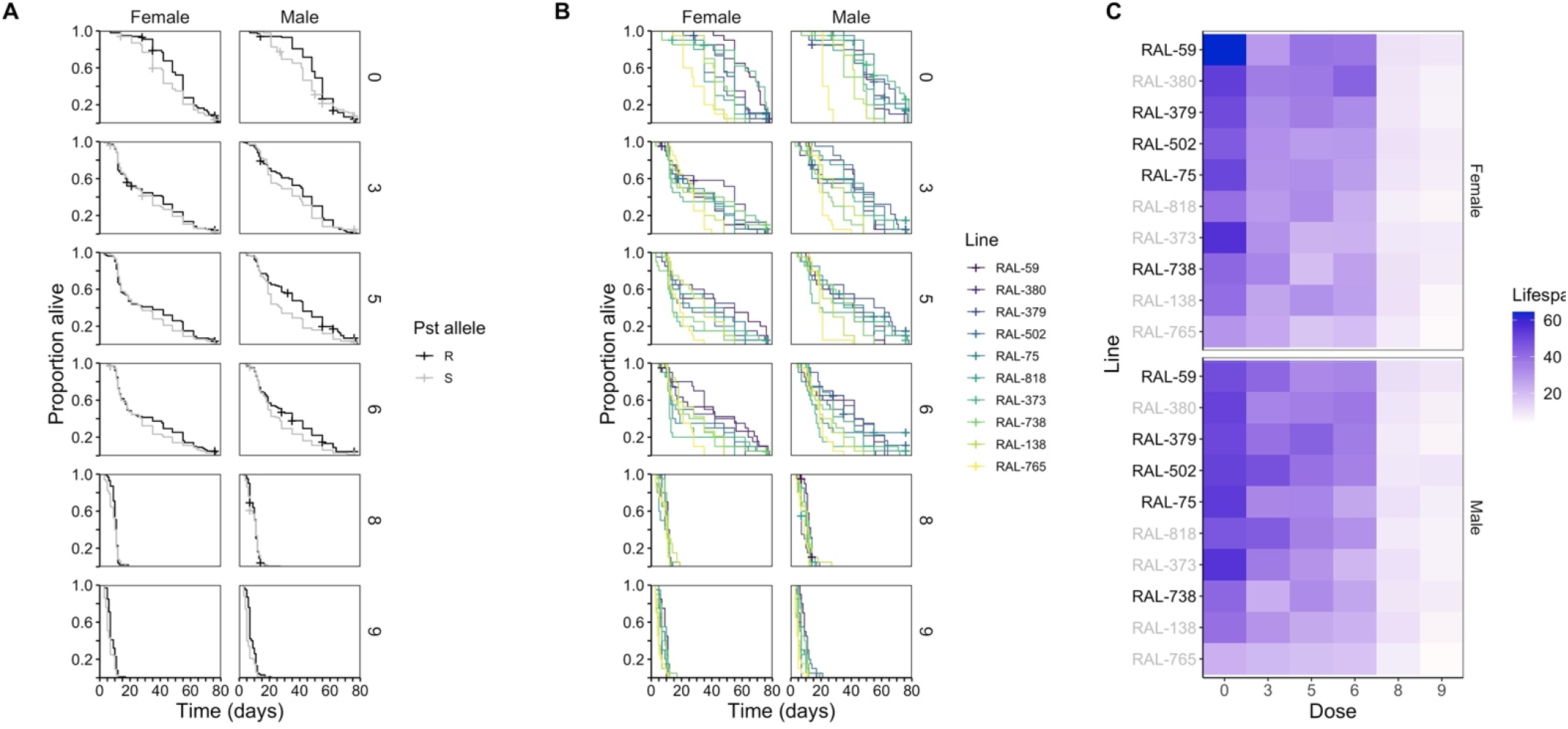
Effects of *pst*, genetic variation, sex and virus dose on survival up to 78 days post infection. Flies were sham treated (Dose 0) or infected with one of five doses (10^3^, 10^5^, 10^6^, 10^8^, 10^9^) of *Drosophila* C Virus. (A) Survival in resistant (R) and susceptible (S) line types. (R lines: RAL-59, RAL-75, RAL-379, RAL-502, RAL-738; S lines: RAL-138, RAL-373, RAL-380, RAL-765, RAL-818). Uninfected resistant lines have a survival advantage in comparison to susceptible lines. Survival tends to improve later in life at low to intermediate infection intensities, but this effect is nearly absent at high DCV doses. (B) Each Kaplan-Meier curve represents the cumulative survival of 20 individuals. Viral dose is logged for ease of interpretation. (C) Heatmaps showing mean lifespan for female (top) and male (bottom) flies, where DGRP lines are arranged according to mean total survival time of males and females. There were differential effects of both line and dose and line and sex on survival after viral infection (B) and (C). R lines are shown in black and S lines are shown in grey. For statistics, see Table 1.

### pastrel is associated with variation in the ability to control viral titres

While *pastrel* has been previously associated with variation in survival following systemic DCV infection, it is not known if *pastrel* acts by improving viral clearance, or if flies carrying the resistant (R) alleles are instead better able to tolerate high viral titres. To test this, we quantified resistance as the rate at which viral titers increased with increasing doses of viral inoculum. This allows a more complete measure of viral clearance for each fly line and sex across several orders of magnitude of viral titre, where a shallow slope indicates the ability to control viral growth even at higher doses, while a steep positive slope suggests that flies lose the ability to control viral growth when exposed to very high doses of DCV. Overall, male and female flies with a resistant (R) *pastrel* allele had significantly lower viral titres compared to susceptible (S) lines (Figure 2A Table 2, Model 2b, *pst* allele: p = 0.003), indicating that *pastrel* explains at least some of the variation in viral titres. For all lines and in both sexes, exposure to higher concentrations of DCV resulted in higher viral titres measured 3 days post infection. However, the magnitude of this increase across DCV doses varied among lines (Figure 2B and 2C, Table 2, Model 2B, Line x Dose: p = 0.0009).

**Table 2.** The effects of DCV dose, sex, and *pst* or DGRP line on resistance. Values in bold are statistically significant.

| Response | Model # | Predictor | Df | F | P-value |
| --- | --- | --- | --- | --- | --- |
| Titre | 2a | Dose | 1 | 417.904 | **<0.0001** |
|  |  | *pst* allele | 1 | 8.897 | **0.003** |
|  |  | Sex | 1 | 0.004 | 0.953 |
| | | Dose × *pst* allele | 1 | 6.751 | **0.0096** |
|  | 2b | Dose | 1 | 77.931 | **<0.0001** |
|  |  | Line | 9 | 4.259 | **<0.0001** |
|  |  | Sex | 1 | 0.004 | 0.952 |
| | | Dose × Line | 9 | 3.626 | **0.0002** |

**Figure 2.**
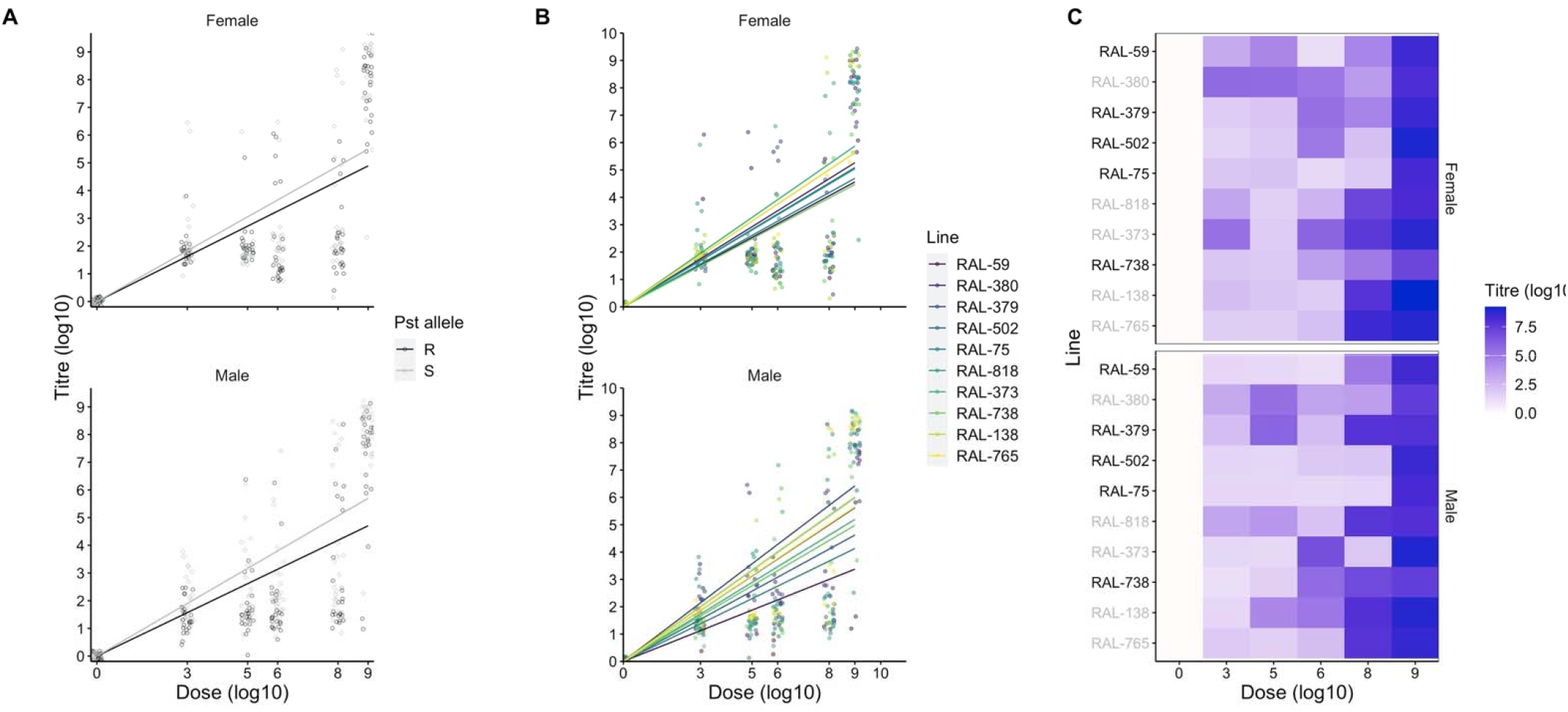
*D. melanogaster* resistance to DCV. (A) DCV titre in R and S DGRP lines, measured three days post infection (3 DPI). R lines: RAL-59, RAL-75, RAL-379, RAL-502, RAL-738; S lines: RAL-138, RAL-373, RAL-380, RAL-, RAL-765, RAL-818. DCV titre is generally lower in resistant DGRP lines. (B) Viral titre measured at 3 DPI differs as a function of sex and line and increase as dose increases. Each data point (n = 5, Line × Sex × Dose) represents the viral titre from a single fly. Values are plotted on log_10_ transformed x– and y– axes. (C) Variation in mean titre for each level of line and dose. Titre is logged for clarity. R lines are shown in black and S lines are shown in grey. For statistics, see Table 2.

### Mortality tolerance to DCV is genetically variable

Since we established that dose was a good indicator of viral load (Figure 2), we used dose as a covariate and a proxy for viral titre in our tolerance models. First, we examined the effect of *pst* on mortality tolerance and found that flies carrying the R allele tended to maintain higher survival over the range of tested doses (higher intercept in Figure 3a; Table 3, Model 3a; *pst* allele: p<0.0001) but we did not detect an effect of *pst* on mortality tolerance (similar definite integrals, when accounting for differences in the intercept). When analysing how survival changes with increasing concentrations of viral challenge, we observed that there was a quadratic relationship between genotype and dose and found that mortality tolerance to DCV was genetically variable (Figure 3B and 3C; Table 3, Model 3b; Dose^2^ × Line: p <0.0001). In order to examine differences in tolerance among lines, the y-intercept of each function was standardized at 0 to account for differences in general vigor, *e*.*g*. Raberg et al 2009, before integration. Here, a small negative integral value (*e*.*g*., RAL-765) indicates a small change in mortality across the tested doses (high tolerance), whereas a large negative integral value (*e*.*g*., RAL-373) indicates large changes in mortality across several orders of magnitude of viral exposure (lower tolerance) (Figure 3C).

**Figure 3.**
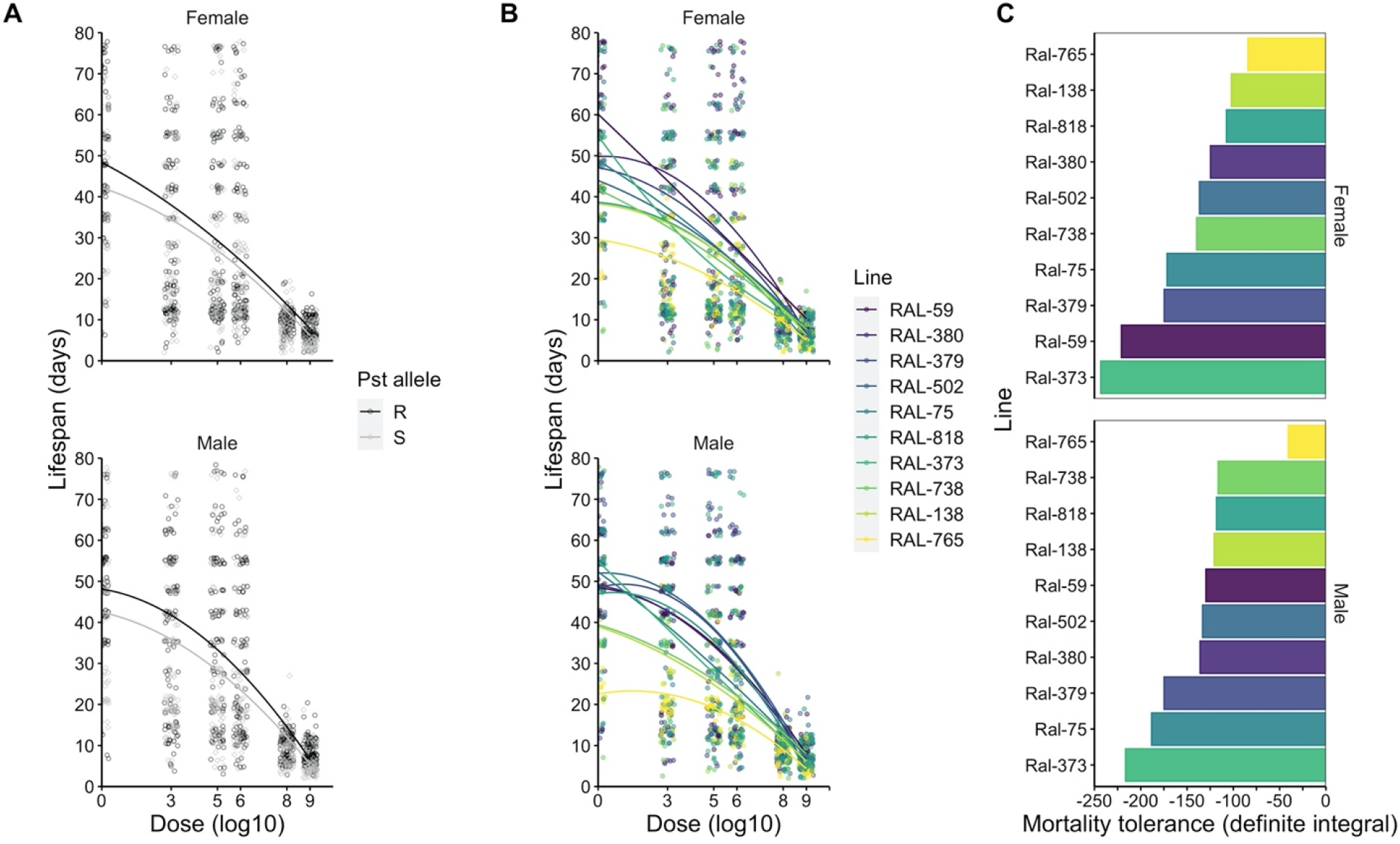
Mortality tolerance in DCV infected flies shows evidence of genetic variation and non-linearity. (A) Lifespan in resistant (R) and susceptible (S) DGRP lines. Resistant lines tend to live longer than susceptible lines and are equally tolerant to DCV infection. R lines: RAL-59, RAL-75, RAL-379, RAL-502, RAL-738; S lines: RAL-138, RAL-373, RAL-380, RAL-, RAL-765, RAL-818. (B) Reaction norms are plotted for each line and split by sex. We use dose in place of titre (*i*.*e*. Lefevre et al 2011, Vale and Gupta 2017) to estimate variation in tolerance. (C) Integrals for each DGRP line, split by sex. The y-intercept of each function was standardized at 0 to account for differences in general vigor, *e*.*g*. [1], before integration. Bars are ordered from least tolerant (Ral-373) to most tolerant (Ral-765).

**Table 3.** The effects of DCV dose and *pst* or DGRP line on mortality tolerance and fecundity tolerance. Values in bold are statistically significant. Models 3a and 3b tested for differences in mortality tolerance between *pst* alleles or among DGRP lines (indicated by a statistically significant interaction with dose or dose^2^). Models 4a and 4b tested for differences in fecundity tolerance between *pst* alleles or among DGRP lines.

| Response | Model # | Predictor | Df | $\chi^2$ | P-value |
| --- | --- | --- | --- | --- | --- |
| Lifespan | 3a | Dose | 1 | 38.227 | **<0.0001** |
|  |  | Dose<sup>2</sup> | 1 | 510.012 | **<0.0001** |
|  |  | pst allele | 1 | 38.6731 | **<0.0001** |
|  |  | Sex | 1 | 2.7915 | 0.09477 |
|  | 3b | Dose | 1 | 1.429 | 0.232 |
|  |  | Dose<sup>2</sup> | 1 | 45.451 | **<0.0001** |
|  |  | Line | 9 | 55.658 | **<0.0001** |
|  |  | Sex | 1 | 0.367 | 0.545 |
| | | Dose × Line | 9 | 33.009 | **0.00013** |
| | | Dose<sup>2</sup> × Line | 9 | 36.088 | **<0.0001** |
| | | Line × Sex | 9 | 20.872 | **0.013** |
| Cumulative fecundity | 4a | Dose | 1 | 5.5437 | **0.0186** |
|  |  | Pst allele | 1 | 97.5767 | **<0.0001** |
|  | 4b | Dose | 1 | 2.1671 | 0.141 |
|  |  | Line | 9 | 88.8993 | **<0.0001** |
| | | Dose × Line | 9 | 16.9953 | **0.0488** |

### Fecundity tolerance of DCV shows evidence of genetic variation

Hosts may tolerate an infection by limiting its negative effects not only on survival but also on reproduction (known as fecundity or sterility tolerance) [7,12,16,20,55,56] so we asked if females from the ten DGRP lines showed variation in fecundity tolerance to DCV. We therefore measured cumulative fecundity (adult offspring production) in single flies over a 28-day period and then quantified fecundity tolerance as the ability to maintain reproduction for increasing viral doses. When accounting for differences in infected lifespan, females with the resistant (R) *pst* allele tended to have more offspring than females with the susceptible (S) allele, (Figure 4A, Table 3; Model 4a, *pst* allele: p < 0.0001). This effect occurred regardless of infection status and R and S lines were equally tolerant, indicated by the similar slopes. The fecundity data further suggest that the R allele is associated with improved reproductive fitness even in the absence of infection (Figure 4A, dose 0).

**Figure 4.**
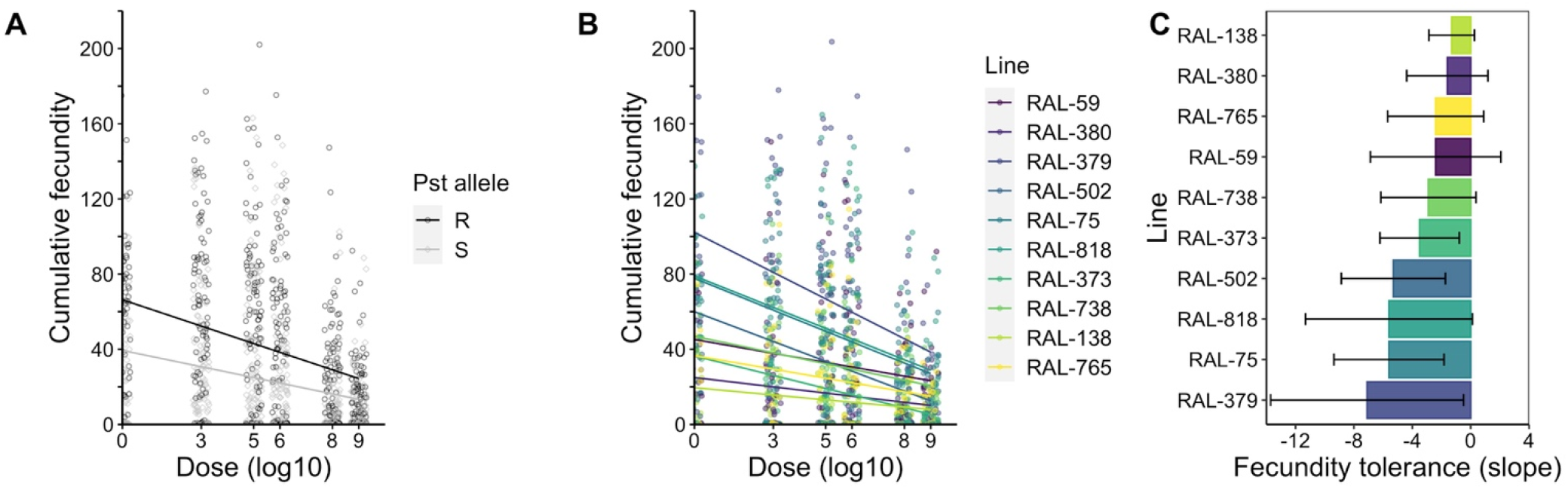
DCV infected DGRP lines show evidence of genetic variation in fecundity tolerance. (A) Cumulative fecundity in R and S DGRP lines. Susceptible lines have fewer offspring than resistant lines regardless of infection status but are equally as tolerant as R lines (similar slopes). (R lines: RAL-59, RAL-75, RAL-138, RAL-373, RAL-379; S lines: RAL-380, RAL-502, RAL-738, RAL-765, RAL-818). (B) Reaction norms are plotted for each DGRP line. Each data point represents the cumulative fecundity of a single fly during its lifetime. (C) Slopes ± SE of reaction norms plotted in (B). Bars represent the fecundity tolerance of each DGRP line. Lines are ordered from the least tolerant (RAL-379) to most tolerant (RAL-138). For statistics see Table 3.

In contrast to mortality tolerance, here the relationship between dose and fecundity was linear and we observed significant differences between lines in the slopes of these linear relationships (Figure 4B, Table 3, Model 4b; Dose × Line: p = 0.0488), although we note that this effect was only marginally significant. To quantify the extent of this decline, we used the slope for each line, where a shallow slope indicates a small change in fecundity across several orders of magnitude of DCV exposure (Figure 4C, *e*.*g*., RAL-138, RAL-380), while steep negative slopes indicate large changes in fecundity with increasing DCV dose, suggesting low fecundity tolerance (Figure 4C, *e*.*g*., RAL-379).

### No evidence of trade-offs between resistance and tolerance

After observing genetic variation in mortality tolerance and fecundity tolerance to DCV, we asked if there was a trade-off between resistance and tolerance. The two strategies are often assumed to exist along a continuum [12,13], but we did not find evidence of such a trade-off (Supplementary Figures S1 and S2). We wondered if we could detect a trade-off between fecundity tolerance and mortality tolerance, as might be expected if investing in fecundity comes at a trade-off with investing in immunity and/or lifespan [57,58]. However, we did not find any evidence of a trade-off between mortality tolerance and fecundity tolerance (Supplementary Figure S3). Overall, our data suggests that the ability to resist or tolerate DCV infection is decoupled in *D. melanogaster*.

### pastrel affects upd3 expression in the absence of infection and G9a expression in infected lines

In a separate experiment, we examined *G9a* and *upd3* expression in males and females infected with a viral concentration of 10^7^ DCV IU ml^-1^. We chose *G9a* because it has been shown to mediate tolerance to DCV infection by regulating the JAK-STAT response[43,44], whereas *upd3* encodes a cytokine-like protein and is the main JAK-STAT ligand induced in response to viral challenge [59]. We reasoned that their expression may explain some variation in disease tolerance and resistance to DCV infection in the ten DGRP lines (Figures 1-3). *pastrel* status was not associated with baseline *G9a* expression in uninfected flies (Figure 5A, Table 4, Model 5a) but we found that *G9a* expression in infected flies was lower in flies carrying a resistant (R) allele versus those carrying a susceptible (S) allele (Figure 5B, Table 4, Model 6a, *pst* allele: p = 0.0007). Baseline *upd3* expression was lower in the S lines (Figure 5C, Table 4, Model 7a, *Pst* allele: p 0.0006) but infected flies showed similar levels of *upd3* expression regardless of their *pastrel* allele (Figure 5D, Table 4, Model 8a).

**Table 4.**
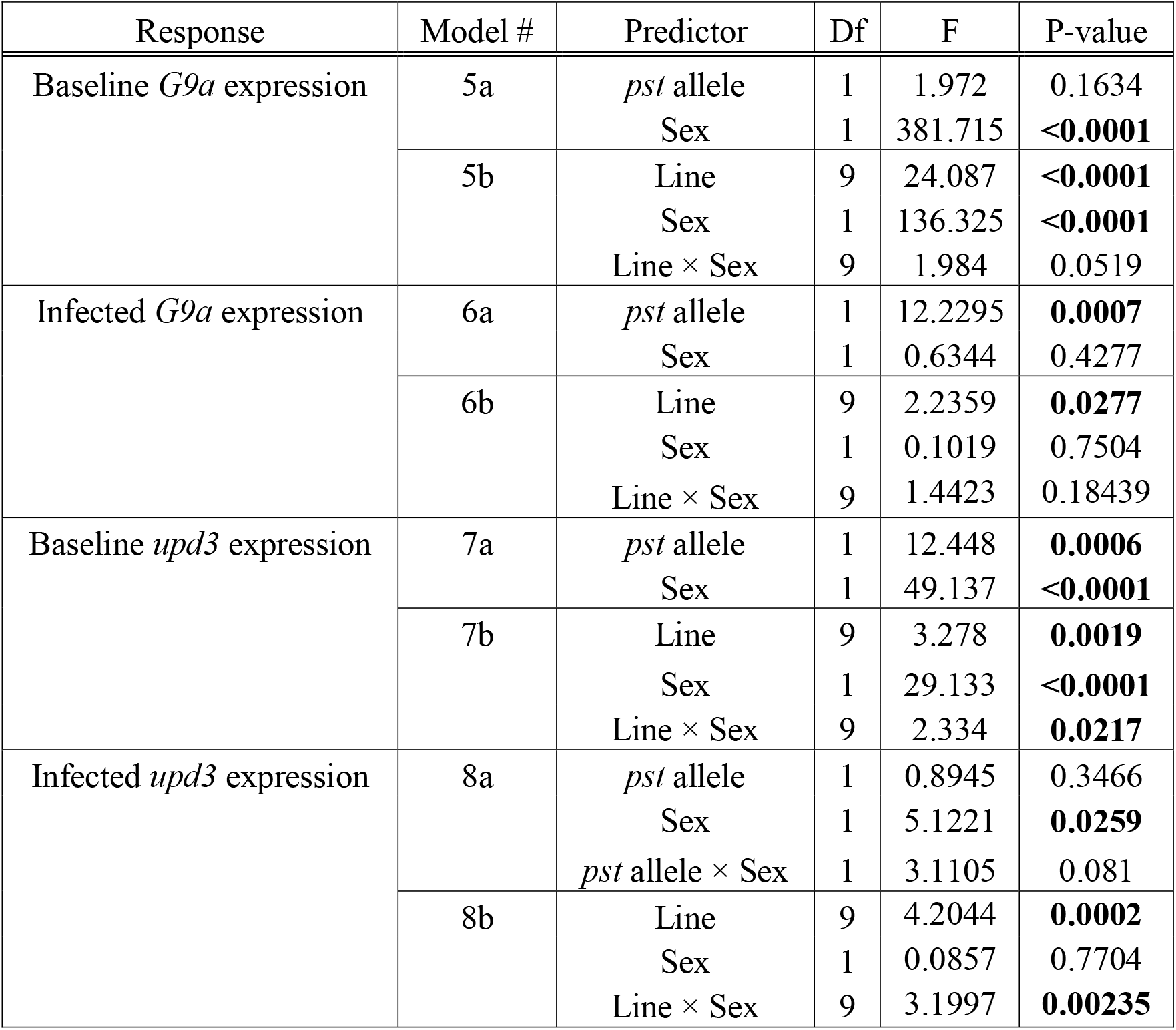
The effects of Sex and *pst* or DGRP line on baseline (relative to rp49) and infected *G9a* or *upd3* expression. Values in bold are statistically significant.

**Figure 5.**
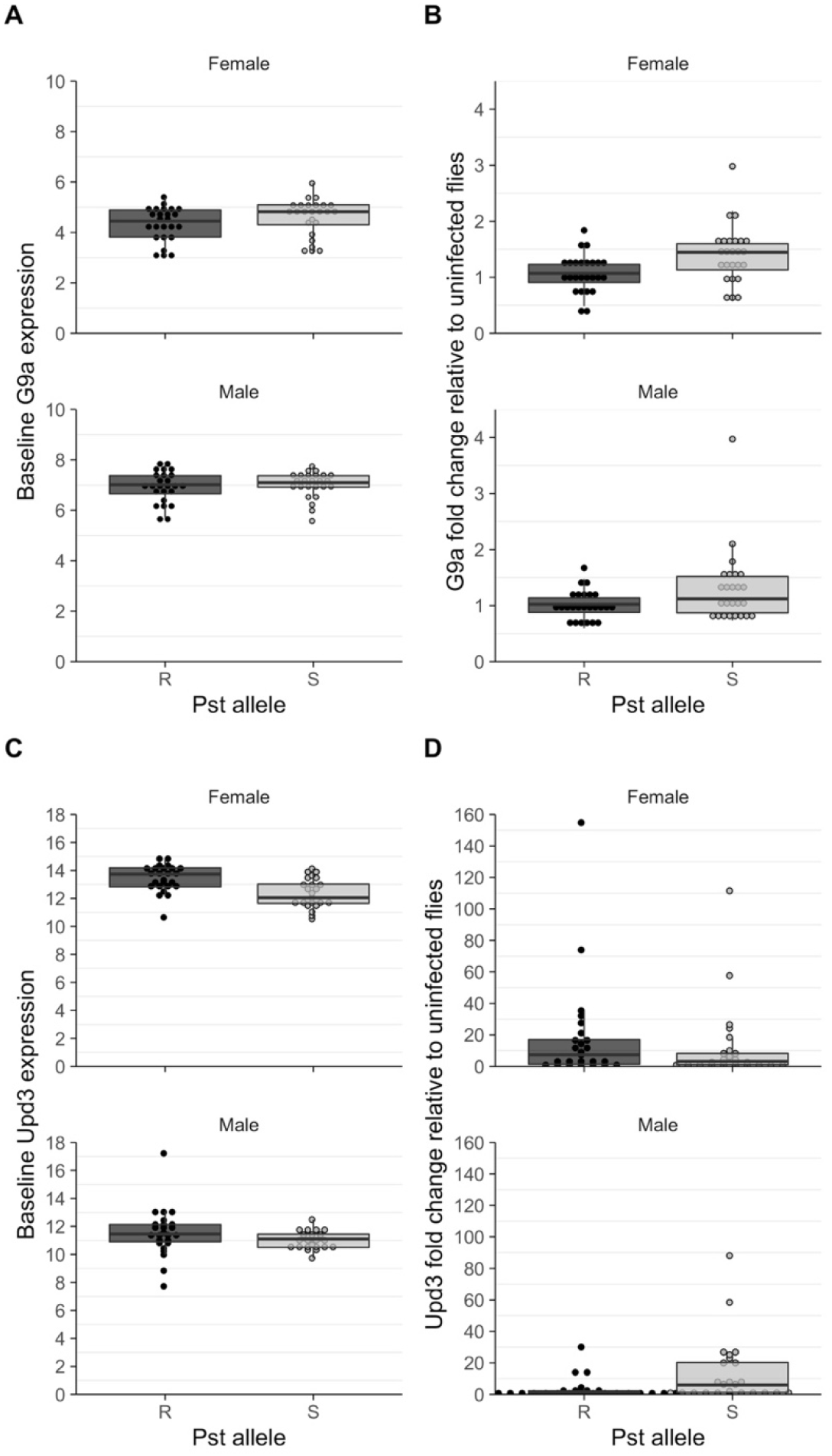
*pst* has differential effects on gene expression between uninfected and infected flies.(A) *G9a* expression relative to *rp49* in the absence of infection is not significantly affected by *pst*. (B) Infected flies *G9a* expression is higher in (S) susceptible DGRP lines but is unaffected by sex. (C) *upd3* expression relative to *rp49* is higher in (R) resistant DGRP lines and tends to be lower in males. (D) Sex affects infected expression of *upd3*. R lines: RAL-59, RAL-75, RAL-379, RAL-502, RAL-738; S lines: RAL-138, RAL-373, RAL-380, RAL-, RAL-765, RAL-818. For statistics see Table 4.

### Genetic variation in the expression of upd3 and G9a does not explain variation in resistance or tolerance

Examining gene expression across all ten lines, we found evidence of genetic variation in *G9a* expression (Figure S4; Table 4; Model 5b, Line: p < 0.0001), and females tended to have lower baseline expression compared with males (Figure S4; Table 4; Model 5b, Sex: p < 0.0001). We found differential effects of sex and line on uninfected *upd3* expression (Figure S4; Table 4; Model 7b; Line × Sex: p = 0.022). In infected flies, *G9a* expression varied between fly lines (Figure S5, Table 4, Model 6b; Line: p = 0.028), and males and females differed in their expression of *upd3* following infection, with males showing generally lower *upd3* expression, although the magnitude of these sex differences varied between DGRP lines (Figure S5; Table 4; Model 8b; Sex x Line: p = 0.002). While both the baseline and infected gene expression differed among fly lines for *G9a* and *upd3*, we did not detect a significant correlation between the expression of either gene and resistance to DCV, mortality tolerance, or fecundity tolerance (Supplementary Figures S6-S17).

### Discussion

We found evidence of genetic variation in disease tolerance in *D. melanogaster* during systemic infection with DCV, measured both as the ability to maintain survival and reproduction, across a wide range of concentrations of viral challenge. We also confirmed results that the viral restriction factor *pastrel* increases fly survival by reducing viral titers, and we further uncovered previously undescribed effects of *pastrel* on general fly vigor in the absence of infection, and it effects on the expression of the JAK-STAT ligand *upd3* and the epigenetic regulator of JAK-STAT, *G9a*.

### pastrel affects host vigor in the absence of infection

The restriction factor *pastrel* has been previously shown to explain most of the variance in fly mortality following systemic DCV infection [22]. Our data confirm these effects, and further confirm that *pst*-mediated increase in fly survival is mainly due to its effects on suppressing DCV titres, which is consistent with its proposed role as a viral restriction factor [29]. The resistant *pst* allele results from a nonsynonymous substitution (A/G; Threonine → Alanine) in the coding region of the gene [29]. The susceptible allele is ancestral and has been shown to play some part in antiviral defense, as overexpression of the allele improves survival after DCV infection and knockdown of the allele makes flies more susceptible to infection. The resulting amino acid substitution is therefore an improvement on an already existing antiviral defense [29].

However, our data also suggest that the effects of *pastrel* extend beyond viral clearance, and in the case of the resistant (R) allele, *pastrel* was associated with a general improvement in fly reproduction and lifespan, even in the absence of infection. To our knowledge, this is the first study to demonstrate *pastrel’s* effects on general vigor. This result is somewhat surprising, because we might expect a mutation that confers antiviral protection to trade-off against other life-history traits [60]. Indeed, in previous work, sham infected control flies that over expressed the S allele tended to live longer than those that over expressed the R allele, suggesting that overexpression of R comes with costs [29]. That study also found natural variation in *pst* gene expression and that its expression is associated with improved survival outcomes after DCV infection, but it is unclear if this is also associated with improved vigor in the absence of infection. Likewise, in a separate study where flies were selected for survival to DCV, *pastrel* was also identified as being involved in adaptation to DCV, with no apparent detrimental effects on egg viability, reproductive output or developmental time [61,62]. Our study confirms that the R allele does not seem to carry costs, but is associated with fitness benefits in the absence of infection. Taken together, it is therefore puzzling why the R allele has not risen to fixation, and why S alleles are maintained in the population. It seems likely that the R allele may come with hidden costs that are not manifested under *ad libitum* laboratory conditions. For example, dietary manipulation can sometimes uncover the costs associated with immunity [16,55,60].

### pastrel controls resistance to DCV

While previous studies established that ‘susceptibility’ to DCV is controlled by *pastrel*, those studies did not directly assay viral loads in resistant versus susceptible natural variants but based their classification on survival data from the DGRP or titre data from knockdown and over expression experiments. These confirmed that the *pastrel* gene confers resistance – viral titres were higher in knockdown flies versus controls and overexpression of both S and R alleles increased resistance – but, crucially, they do not establish whether *pastrel* underlies variation in viral titre in natural fly populations [22,29]. Given these results, there were two possibilities: 1) the R allele confers resistance by controlling viral titres or 2) the R allele confers tolerance to DCV by maintaining survival or reducing damage in the face of infection. Our results support the first possibility that the R allele promotes resistance, demonstrated by lower viral titres in DGRP lines carrying the R allele, and that this protective effect was present in males and female flies, across several orders of magnitude of viral challenge.

### Genetic variation in mortality tolerance and fecundity tolerance

Fly genetic background affected the ability of flies to tolerate DCV infection, both when tolerating the mortality caused by infection, and by maintaining fecundity at low and intermediate viral challenge doses. Previous theoretical work showed that variation in fecundity tolerance is more likely to occur if it comes at a cost to host lifespan or another life history trait [12]. Although we did not observe a trade-off with survival or mortality tolerance in our system, it is possible that fecundity tolerance comes at a cost to another trait that we did not measure. Evidence for genetic variation in both mortality and fecundity tolerance phenotypes is widespread throughout the animal kingdom (reviewed in [4,5]), reinforcing the idea that disease tolerance is an important defence strategy in response to a range of pathogens. It is notable however, that most experimental studies examining genetic variation in disease tolerance have rarely measured it in the context of viral infections [63]. Our work is, to our knowledge, the first to describe genetic variation in both mortality and fecundity tolerance of a viral infection.

### Linear and non-linear changes in health

The majority of tolerance experiments often assume a linear relationship between pathogen load and host health (or other fitness trait), but there is no reason to assume that health should decrease at a constant rate in relation to pathogen burden [1,44,64–66]. We show that some genotypes maintain their health (measured as lifespan) at a low and intermediate DCV doses, whereas health declines rapidly in others. Similar nonlinear relationships between pathogen load and health occur over the course of natural HIV infection in humans [65], in blue tits (*Cyanistes caerulus*) infected with the blood parasite, *Haemoproteus majoris* [64], and in *Drosophila melanogaster* infected with *Listeria monocytogenes* [66] or DCV [44]. In contrast, we found that the relationship between cumulative fecundity and viral dose was best explained by a linear relationship although previous studies on DCV’s effects on fecundity note that offspring production tends to increase at low or intermediate viral doses [26].

### No sex differences in tolerance or resistance to DCV

Sexual dimorphism in immunity is widespread across metazoans, and to a large extent has frequently been overlooked in experimental studies of infection [67–69]. The sexes can differ in optimal immune investment and allocate resources to different areas of the immune response [15,70–72]. In general, females tend to be more immunocompetent than males because they improve their fitness by increasing investment in immune defence, known as Bateman’s principle [70,71,73]. In systems where resistance and tolerance are negatively correlated as shown in malaria infected mice [13], one sex may invest more into resistance, while the other may invest in tolerance. Sex differences in disease tolerance are also predicted to have qualitatively different consequences for pathogen evolutionary trajectories [72].

It was therefore an explicit aim of the present study to quantify sex differences in lifespan, resistance and disease tolerance following DCV infection, to examine potential sexual dimorphism in disease tolerance. However, we were surprised to find that fly sex contributed little to the variation in the disease phenotypes we investigated, particularly viral titers or mortality tolerance. This contrasts with some results from disease tolerance in other host-pathogen systems where sexual dimorphism in tolerance has been observed (reviewed in [72]). For example, males infected with *P. aeruginosa* were more tolerant and resistant than females, with evidence of sexual antagonism for tolerance, indicated by a negative genetic intersexual correlation[15]. By contrast, Gupta and Vale [26] noted that *D. melanogaster* males are more susceptible than females to systemic DCV infection, while no difference between males and females was detected in tolerance of HIV [65]. It is therefore difficult to make generalizations concerning disease outcomes between the sexes, which will depend on the specific host and pathogen species, particularly as the expression of many infection-related traits is often the outcome of complex interactions between host sex, genetic background, and mating status [30]. What is clearer is that work reporting sex-specific infection outcomes are less common than is desirable, especially regarding disease tolerance phenotypes.

### pastrel is associated with changes in pre- and post-infection gene expression

Given previous work [43,44], we expected that *G9a* and *upd3* expression would correlate with disease tolerance and explain some of the phenotypic variation we see among DGRP lines. Although we observed differential effects of genetic background and sex in gene expression, this appeared to be independent of disease tolerance phenotypes. We note that *pastrel* was associated with differences in baseline *upd3* expression as well as infected *G9a* expression. Baseline *upd3* expression was lower in susceptible lines, suggesting that expression levels prior to infection may dictate the speed or strength of the antiviral immune response. Differences in baseline gene expression have been shown to affect chronic disease outcomes (*e*.*g*., rheumatoid arthritis, multiple sclerosis, lung cancer, autoimmune diseases) [68,74–76], so we suggest that basal expression levels may be important predictors of resistance and tolerance. Similarly, infected *G9a* expression was higher in susceptible lines, which may point to differences in the damage control response which we were unable to detect as a tolerance phenotype in our experiments. In fact, it is possible that *G9a* expression may not be directly related to DCV infection at all, as recent work has highlighted the likely role of this methyltransferase as a master regulator of metabolic homeostasis and tolerance to a variety of biotic and abiotic stressors in many different species [77].

### Concluding remarks

In summary, we describe genetic variation in disease tolerance in *Drosophila* following systemic DCV infection, in males and females, across a range of infectious challenges spanning several orders of magnitude. Further, we find that the *pastrel* gene is associated with general vigor in the absence of infection and confirm its role in reducing DCV titres during infection. This work offers, to our knowledge, one of the first descriptions of genetic variation in mortality and fecundity tolerance in a viral infection of invertebrates, adding to the growing effort to describe the causes of host heterogeneity in order to predict the consequences of this heterogeneity for pathogen spread and evolution [78–80].

## Acknowledgements

We are grateful to H. Borthwick and H. Cowan for help with media preparation and members of the Obbard lab for advice on DCV infection culture.

