## Appendix S1 for "The restriction factor *pastrel* is associated with host vigor, viral titer, and variation in disease tolerance during Drosophila C Virus infection"

***Statistical analyses***

**Model 1: Effect of dose, sex, and *pst* or genotype on survival**

Survival over the experimental period was analyzed using Cox mixed effects models [1] with day of death as the response variable and experimental block as a random intercept. Flies that were still alive at the end of the experiment were included as censored cases. Model 1a compared survival between *pst* alleles and Model 1b compared survival among genotypes. Dose was included as a covariate in both models and was log transformed. We started with the full model in both cases and evaluated reduced model likelihoods using model selection criteria. Both models were additionally tested without random effects to test whether they fulfilled the assumptions of proportional hazards over time using the cox.zph function [2]. Both models did not fulfill the assumptions of proportional hazards, so we used a survival regression (survreg function in the survival package [3]) to check that these models gave qualitatively similar results to the Cox mixed effects models. The models explaining significant variance in survival after model reduction are detailed below.

Model 1a: Survival ~ Dose + *pst* allele + Sex + (1|Block)

Model 1b: Survival ~ Dose + Line + Sex + Dose × Line + Line × Sex + (1|Block)

**Model 2: Effect of dose, sex and *pst* or genotype on viral titre**

We examined resistance to DCV using generalized linear models with gamma errors. Both titre and dose were log transformed. We note that our dataset contained some false positives in our uninfected control groups which was likely due to over sensitivity in our detection threshold. Therefore, we analyzed the titre in two ways: using the false positives and by changing the false positives to zeros. Our results were virtually identical in both cases, although we note that the inclusion of zero values caused some problems with the variance in the models. However, we chose to include model results with the zero values here and in the figures for ease of interpretation. We note that our dataset shows evidence of nonlinearity but logging the response variable and the covariate should have accounted for this. Models were reduced using model selection criteria which resulted in the following final models:

Model 2a: Titre ~ Dose + *pst* allele + Sex + Dose × *pst* allele

Model 2b: Titre ~ Dose + Line + Sex + Dose + Dose × Line

**Model 3: Effect of dose, sex, and pst or genotype on mortality tolerance**

We analyzed mortality tolerance using generalized linear mixed models with negative binomial errors and a quadratic parameterization (nbinom2 in the glmmTMB package [4]) to account for overdispersion. We used lifespan as a response variable and included log transformed dose and dose^2^ as covariates. Any significant interaction between the covariates and our fixed effects of *pst* allele, sex or genotype (line) indicates significant changes in mortality tolerance. We included experimental block as a random intercept in our models. Models were reduced using model selection criteria. This resulted in the following models:

Model 3a: Lifespan ~ Dose + Dose^2^ + *pst* allele + Sex + (1|Block)

Model 3b: Lifespan ~ Dose + Dose^2^ + Line + Sex + Dose × Line + Dose^2^ × Line + Line × Sex + (1|Block)

**Model 4: Effect of dose and *pst* or genotype on fecundity tolerance**

Models 4a and 4b evaluated changes in fecundity tolerance between *pst* alleles (4a) or among the 10 tested DGRP lines (4b) using negative binomial generalized linear mixed models with a linear parameterization (nbinom1 in the glmmTMB package [4]) to account for overdispersion and zero inflation. We used total reproductive days as an offset term to account for differences in reproductive effort among females and included experimental block as a random intercept in both models. We started out with the full model in both cases, which included Dose^2^ but we did not detect evidence of nonlinearity, so it was removed from both models. Dose was log transformed in both models. Any significant interaction between the covariate and our fixed effects of *pst* allele or genotype (line) indicates significant changes in fecundity tolerance. Models were further reduced using model selection criteria. After reduction, the models explaining significant variation in fecundity tolerance were as follows:

Model 4a: Cumulative fecundity ~ Dose + *pst* allele + offset(Reproductive days) + (1|Block)

Model 4b: Cumulative fecundity ~ Dose + Line + Dose × Line + offset(Reproductive days) + (1|Block)

**Models 5 – 8: Effect of sex and *pst* or genotype on gene expression**

Models 5-8 evaluated changes in baseline gene expression and infected gene expression between *pst* alleles, among lines, and between the sexes using linear models. The residuals were reasonably normally distributed. We started with the full models in all cases and evaluated reduced model likelihoods using model selection criteria. The final models are detailed below:

Model 5a: Baseline G9a expression ~ *pst* allele + Sex

Model 5b: Baseline G9a expression ~ Line + Sex + Line × Sex

Model 6a: Infected G9a expression ~ *pst* allele + Sex

Model 6b: Infected G9a expression ~ Line + Sex + Line × Sex

Model 7a: Baseline upd3 expression ~ *pst* allele + Sex

Model 7b: Baseline upd3 expression ~ Line + Sex + Line × Sex

Model 8a: Infected upd3 expression ~ *pst* allele + Sex + pst allele × Sex

Model 8b: Infected upd3 expression ~ Line + Sex + Line × Sex
