## Supplementary Table for "The restriction factor *pastrel* is associated with host vigor, viral titer, and variation in disease tolerance during Drosophila C Virus infection"

**Supplementary tables**

**Table S1: qPCR primers**

| **Gene Target** | **Primers** |
| --- | --- |
| *G9a* | Forward_5’ CGTAGCGTTCGTTTAGGCAA 3’ |
|  | Reverse_5’ TTGGGTGGGGTCTTCTCATC 3’ |
| *Upd3* | Forward_5’ AGTGAGCACCAAGACTCTGG 3’ |
|  | Reverse_5’ GTGGCGAAGGTTCAACTGTT 3’ |
| *rp49* | Forward_5’ ATGCTAAGCTGTCGCACAAATG 3’ |
|  | Reverse_5’ GTTCGATCCGTAACCGATGT 3’ |

**Table S2: Model 1a**

| Fixed coefficients | |  |  |  |  |
| --- | --- | --- | --- | --- | --- |
|  | **coef** | **exp(coef)** | **se(coef)** | **z** | **p** |
| **Line.typeS** | 0.222 | 1.248588 | 0.04188 | 5.3 | 1.20E-07 |
| **log10(Dose)** | 0.309 | 1.36193 | 0.01003 | 30.8 | 0.00E+00 |
| **SexMale** | -0.07 | 0.93179 | 0.04167 | -1.7 | 9.00E-02 |

| Random effects | |  |  |
| --- | --- | --- | --- |
| **Group** | **Variable** | **Std Dev** | **Variance** |
| Block | Intercept | 0.177191 | 0.0314 |

**Table S3:** **Model 1b**

| Fixed coefficients |  |  |  |  |  |
| --- | --- | --- | --- | --- | --- |
|  | **coef** | **exp(coef)** | **se(coef)** | **z** | **p** |
| **LineRAL-380** | -0.0962439 | 0.9082425 | 0.24482865 | -0.39 | 6.90E-01 |
| **LineRAL-379** | -0.1489701 | 0.8615949 | 0.25673422 | -0.58 | 5.60E-01 |
| **LineRAL-502** | 0.28781072 | 1.3335049 | 0.2439329 | 1.18 | 2.40E-01 |
| **LineRAL-75** | 0.1569309 | 1.1699148 | 0.24252031 | 0.65 | 5.20E-01 |
| **LineRAL-818** | 0.25660793 | 1.2925383 | 0.25061757 | 1.02 | 3.10E-01 |
| **LineRAL-373** | -0.0919882 | 0.912116 | 0.23646063 | -0.39 | 7.00E-01 |
| **LineRAL-738** | 0.63867501 | 1.8939697 | 0.23154693 | 2.76 | 5.80E-03 |
| **LineRAL-138** | 0.75462872 | 2.1268217 | 0.23833335 | 3.17 | 1.50E-03 |
| **LineRAL-765** | 1.00017736 | 2.718764 | 0.24316067 | 4.11 | 3.90E-05 |
| **log10(Dose)** | 0.28603911 | 1.3311445 | 0.02725869 | 10.49 | 0.00E+00 |
| **SexMale** | 0.23842346 | 1.2692466 | 0.13234981 | 1.8 | 7.20E-02 |
| **LineRAL-380:log10(Dose)** | 0.02817019 | 1.0285707 | 0.04044759 | 0.7 | 4.90E-01 |
| **LineRAL-379:log10(Dose)** | 0.09198805 | 1.0963517 | 0.04206451 | 2.19 | 2.90E-02 |
| **LineRAL-502:log10(Dose)** | 0.02517322 | 1.0254927 | 0.03927884 | 0.64 | 5.20E-01 |
| **LineRAL-75:log10(Dose)** | 0.03266179 | 1.033201 | 0.03964538 | 0.82 | 4.10E-01 |
| **LineRAL-818:log10(Dose)** | 0.07639437 | 1.0793882 | 0.04104763 | 1.86 | 6.30E-02 |
| **LineRAL-373:log10(Dose)** | 0.0955147 | 1.100225 | 0.0379186 | 2.52 | 1.20E-02 |
| **LineRAL-738:log10(Dose)** | -0.0243577 | 0.9759365 | 0.03759868 | -0.65 | 5.20E-01 |
| **LineRAL-138:log10(Dose)** | -0.0176529 | 0.982502 | 0.03847006 | -0.46 | 6.50E-01 |
| **LineRAL-765:log10(Dose)** | 0.00744067 | 1.0074684 | 0.03964477 | 0.19 | 8.50E-01 |
| **LineRAL-380:SexMale** | -0.0163998 | 0.983734 | 0.18504577 | -0.09 | 9.30E-01 |
| **LineRAL-379:SexMale** | -0.4863178 | 0.6148863 | 0.18810141 | -2.59 | 9.70E-03 |
| **LineRAL-502:SexMale** | -0.647462 | 0.5233724 | 0.18870978 | -3.43 | 6.00E-04 |
| **LineRAL-75:SexMale** | -0.3206193 | 0.7256994 | 0.18820003 | -1.7 | 8.80E-02 |
| **LineRAL-818:SexMale** | -0.6917124 | 0.5007179 | 0.18829321 | -3.67 | 2.40E-04 |
| **LineRAL-373:SexMale** | -0.3258326 | 0.721926 | 0.18644328 | -1.75 | 8.10E-02 |
| **LineRAL-738:SexMale** | -0.1279441 | 0.8799025 | 0.18510629 | -0.69 | 4.90E-01 |
| **LineRAL-138:SexMale** | -0.1750897 | 0.8393817 | 0.18527236 | -0.95 | 3.40E-01 |
| **LineRAL-765:SexMale** | -0.1829289 | 0.8328274 | 0.1849525 | -0.99 | 3.20E-01 |
| Random effects |  |  |  |  |  |
| **Group** | **Variable** | **Std Dev** | **Variance** |  |  |
| Block | Intercept | 0.16967949 | 0.02879113 |  |  |

**Table S4:** **Model 2a**

| Coefficients |  |  |  |  |
| --- | --- | --- | --- | --- |
|  | **Estimate** | **Std. Error** | **t value** | **Pr(>\|t\|)** |
| **(Intercept)** | 1.2927107 | 0.0643372 | 20.093 | < 2e-16 |
| **Line.typeS** | -0.2450524 | 0.0826914 | -2.963 | 0.00316 |
| **log10(Dose)** | -0.1278201 | 0.0075556 | -16.917 | < 2e-16 |
| **SexMale** | 0.0008372 | 0.0141803 | 0.059 | 0.95294 |
| **Line.typeS:log10(Dose)** | 0.0253359 | 0.009796 | 2.586 | 0.00994 |

**Table S5: Model 2b**

| Coefficients: |  |  |  |  |
| --- | --- | --- | --- | --- |
|  | **Estimate** | **Std. Error** | **t value** | **Pr(>\|t\|)** |
| **(Intercept)** | 1.4183837 | 0.1549838 | 9.152 | < 2e-16 |
| **log10(Dose)** | -0.1377242 | 0.0184889 | -7.449 | 3.43E-13 |
| **SexMale** | 0.0008228 | 0.0136123 | 0.06 | 0.95182 |
| **LineRAL-380** | -0.5838426 | 0.1820263 | -3.207 | 0.00141 |
| **LineRAL-379** | -0.4964091 | 0.1848842 | -2.685 | 0.00746 |
| **LineRAL-502** | 0.0598556 | 0.2184155 | 0.274 | 0.78415 |
| **LineRAL-75** | 0.1275953 | 0.2244815 | 0.568 | 0.56998 |
| **LineRAL-818** | -0.5455042 | 0.1836386 | -2.971 | 0.0031 |
| **LineRAL-373** | -0.2341229 | 0.1993022 | -1.175 | 0.24059 |
| **LineRAL-738** | -0.1074435 | 0.2090297 | -0.514 | 0.60744 |
| **LineRAL-138** | -0.2136791 | 0.1996436 | -1.07 | 0.28493 |
| **LineRAL-765** | -0.1566126 | 0.2032489 | -0.771 | 0.44129 |
| **log10(Dose):LineRAL-380** | 0.0610663 | 0.0219188 | 2.786 | 0.00551 |
| **log10(Dose):LineRAL-379** | 0.0485229 | 0.0220913 | 2.196 | 0.02845 |
| **log10(Dose):LineRAL-502** | -0.0118182 | 0.0257708 | -0.459 | 0.6467 |
| **log10(Dose):LineRAL-75** | -0.017565 | 0.0265283 | -0.662 | 0.50816 |
| **log10(Dose):LineRAL-818** | 0.0561689 | 0.0220642 | 2.546 | 0.01116 |
| **log10(Dose):LineRAL-373** | 0.0195308 | 0.0236492 | 0.826 | 0.40923 |
| **log10(Dose):LineRAL-738** | 0.0081323 | 0.0248445 | 0.327 | 0.74354 |
| **log10(Dose):LineRAL-138** | 0.0162125 | 0.0236296 | 0.686 | 0.49292 |
| **log10(Dose):LineRAL-765** | 0.0102038 | 0.0240328 | 0.425 | 0.6713 |

**Table S6: Model 3a**

|  | **Estimate** | **Std. Error** | **z value** | **Pr(>\|z\|)** |
| --- | --- | --- | --- | --- |
| **(Intercept)** | 3.831775 | 0.053445 | 71.7 | < 2e-16 |
| **Line.typeS** | -0.143955 | 0.023149 | -6.22 | 5.01E-10 |
| **SexMale** | 0.038646 | 0.023131 | 1.67 | 0.0948 |
| **log10(Dose)** | 0.076828 | 0.012426 | 6.18 | 6.30E-10 |
| **I((log10(Dose))^2)** | -0.030833 | 0.001365 | -22.58 | < 2e-16 |

**Table S7: Model 3b**

|  | **Estimate** | **Std. Error** | **z value** | **Pr(>\|z\|)** |
| --- | --- | --- | --- | --- |
| **(Intercept)** | 3.9843199 | 0.0919162 | 43.35 | < 2e-16 |
| **LineRAL-380** | -0.1076019 | 0.1173453 | -0.92 | 0.35916 |
| **LineRAL-379** | -0.1901769 | 0.1175854 | -1.62 | 0.1058 |
| **LineRAL-502** | -0.2300382 | 0.1187995 | -1.94 | 0.05282 |
| **LineRAL-75** | -0.102203 | 0.1182263 | -0.86 | 0.38733 |
| **LineRAL-818** | -0.3740642 | 0.1188502 | -3.15 | 0.00165 |
| **LineRAL-373** | -0.019215 | 0.1181528 | -0.16 | 0.87081 |
| **LineRAL-738** | -0.3154471 | 0.1183903 | -2.66 | 0.00771 |
| **LineRAL-138** | -0.3578475 | 0.1186991 | -3.01 | 0.00257 |
| **LineRAL-765** | -0.7126307 | 0.1196948 | -5.95 | 2.62E-09 |
| **SexMale** | -0.0418081 | 0.0690333 | -0.61 | 0.54477 |
| **log10(Dose)** | 0.044139 | 0.0369186 | 1.2 | 0.23186 |
| **I((log10(Dose))^2)** | -0.0261078 | 0.0040551 | -6.44 | 1.21E-10 |
| **LineRAL-380:SexMale** | 0.0092744 | 0.0978932 | 0.09 | 0.92452 |
| **LineRAL-379:SexMale** | 0.1139148 | 0.0982713 | 1.16 | 0.24638 |
| **LineRAL-502:SexMale** | 0.252053 | 0.0980821 | 2.57 | 0.01018 |
| **LineRAL-75:SexMale** | 0.0745447 | 0.098285 | 0.76 | 0.44818 |
| **LineRAL-818:SexMale** | 0.258064 | 0.0986961 | 2.61 | 0.00893 |
| **LineRAL-373:SexMale** | 0.0869415 | 0.0985228 | 0.88 | 0.37753 |
| **LineRAL-738:SexMale** | 0.0404789 | 0.098443 | 0.41 | 0.68093 |
| **LineRAL-138:SexMale** | 0.0070244 | 0.0988412 | 0.07 | 0.94334 |
| **LineRAL-765:SexMale** | -0.0728605 | 0.1003743 | -0.73 | 0.46791 |
| **LineRAL-380:log10(Dose)** | 0.0985742 | 0.052259 | 1.89 | 0.05926 |
| **LineRAL-379:log10(Dose)** | 0.1201338 | 0.0523837 | 2.29 | 0.02183 |
| **LineRAL-502:log10(Dose)** | 0.0283947 | 0.0526884 | 0.54 | 0.58994 |
| **LineRAL-75:log10(Dose)** | -0.0168967 | 0.0525284 | -0.32 | 0.7477 |
| **LineRAL-818:log10(Dose)** | 0.0935741 | 0.0527895 | 1.77 | 0.0763 |
| **LineRAL-373:log10(Dose)** | -0.120313 | 0.0528908 | -2.27 | 0.02292 |
| **LineRAL-738:log10(Dose)** | -0.0079493 | 0.052912 | -0.15 | 0.88058 |
| **LineRAL-138:log10(Dose)** | 0.0498827 | 0.053093 | 0.94 | 0.34746 |
| **LineRAL-765:log10(Dose)** | 0.0805824 | 0.054064 | 1.49 | 0.13609 |
| **LineRAL-380:I((log10(Dose))^2)** | -0.0127973 | 0.0057826 | -2.21 | 0.02689 |
| **LineRAL-379:I((log10(Dose))^2)** | -0.0166796 | 0.0057928 | -2.88 | 0.00398 |
| **LineRAL-502:I((log10(Dose))^2)** | -0.0034112 | 0.0057574 | -0.59 | 0.55352 |
| **LineRAL-75:I((log10(Dose))^2)** | 0.0002123 | 0.0057758 | 0.04 | 0.97068 |
| **LineRAL-818:I((log10(Dose))^2)** | -0.0130505 | 0.0058061 | -2.25 | 0.02459 |
| **LineRAL-373:I((log10(Dose))^2)** | 0.010506 | 0.005795 | 1.81 | 0.06984 |
| **LineRAL-738:I((log10(Dose))^2)** | 0.0026815 | 0.0058017 | 0.46 | 0.64395 |
| **LineRAL-138:I((log10(Dose))^2)** | -0.0056217 | 0.0058647 | -0.96 | 0.33777 |
| **LineRAL-765:I((log10(Dose))^2)** | -0.0069703 | 0.0059598 | -1.17 | 0.24218 |

**Table S8: Model 4a**

| Conditional model: | |  | |  |  |
| --- | --- | --- | --- | --- | --- |
|  | **Estimate** | | **Std. Error** | **z value** | **Pr(>\|z\|)** |
| **(Intercept)** | 1.001759 | | 0.050839 | 19.704 | <2e-16 |
| **Line.typeS** | -0.553034 | | 0.055986 | -9.878 | <2e-16 |
| **log10(Dose)** | 0.021596 | | 0.009172 | 2.355 | 0.0185 |

| Zero-inflation model: | |  |  |  |
| --- | --- | --- | --- | --- |
|  | **Estimate** | **Std. Error** | **z value** | **Pr(>\|z\|)** |
| **(Intercept)** | -4.96123 | 0.34454 | -14.4 | < 2e-16 |
| **Line.typeS** | -0.0123 | 0.17605 | -0.07 | 0.944 |
| **log10(Dose)** | 0.14649 | 0.03207 | 4.568 | 4.92E-06 |

**Table S9: Model 4b**

| Conditional model: |  |  |  |  |
| --- | --- | --- | --- | --- |
|  | **Estimate** | **Std. Error** | **z value** | **Pr(>\|z\|)** |
| **(Intercept)** | 0.795153 | 0.129615 | 6.135 | 8.53E-10 |
| **LineRAL-380** | -0.263576 | 0.208376 | -1.265 | 0.20591 |
| **LineRAL-379** | 0.426571 | 0.162264 | 2.629 | 0.00857 |
| **LineRAL-502** | -0.105298 | 0.178131 | -0.591 | 0.55444 |
| **LineRAL-75** | 0.263842 | 0.162712 | 1.622 | 0.1049 |
| **LineRAL-818** | 0.377012 | 0.166861 | 2.259 | 0.02386 |
| **LineRAL-373** | -0.593858 | 0.198009 | -2.999 | 0.00271 |
| **LineRAL-738** | -0.03148 | 0.179853 | -0.175 | 0.86106 |
| **LineRAL-138** | -1.002237 | 0.221306 | -4.529 | 5.93E-06 |
| **LineRAL-765** | -0.143923 | 0.207302 | -0.694 | 0.48751 |
| **log10(Dose)** | 0.03586 | 0.024359 | 1.472 | 0.14099 |
| **LineRAL-380:log10(Dose)** | -0.125966 | 0.040622 | -3.101 | 0.00193 |
| **LineRAL-379:log10(Dose)** | 0.020584 | 0.030853 | 0.667 | 0.50466 |
| **LineRAL-502:log10(Dose)** | -0.012119 | 0.034733 | -0.349 | 0.72716 |
| **LineRAL-75:log10(Dose)** | -0.013927 | 0.031262 | -0.445 | 0.65597 |
| **LineRAL-818:log10(Dose)** | -0.014912 | 0.032779 | -0.455 | 0.64917 |
| **LineRAL-373:log10(Dose)** | 0.002235 | 0.039796 | 0.056 | 0.95521 |
| **LineRAL-738:log10(Dose)** | -0.019662 | 0.033198 | -0.592 | 0.55368 |
| **LineRAL-138:log10(Dose)** | -0.008079 | 0.040227 | -0.201 | 0.84084 |
| **LineRAL-765:log10(Dose)** | 0.021024 | 0.038817 | 0.542 | 0.58808 |
| Zero-inflation model: |  |  |  |  |
|  | **Estimate** | **Std. Error** | **z value** | **Pr(>\|z\|)** |
| **(Intercept)** | -4.10632 | 0.51632 | -7.953 | 1.82E-15 |
| **LineRAL-380** | 0.60023 | 0.59822 | 1.003 | 0.31569 |
| **LineRAL-379** | -1.62508 | 0.68492 | -2.373 | 0.01766 |
| **LineRAL-502** | -1.86797 | 0.73178 | -2.553 | 0.01069 |
| **LineRAL-75** | -2.15351 | 0.81739 | -2.635 | 0.00842 |
| **LineRAL-818** | -0.9853 | 0.64635 | -1.524 | 0.12741 |
| **LineRAL-373** | -2.72155 | 0.96481 | -2.821 | 0.00479 |
| **LineRAL-738** | -0.60765 | 0.6835 | -0.889 | 0.37399 |
| **LineRAL-138** | -0.99964 | 0.7702 | -1.298 | 0.19433 |
| **LineRAL-765** | 0.16073 | 0.58781 | 0.273 | 0.78451 |
| **log10(Dose)** | 0.12807 | 0.06869 | 1.864 | 0.06228 |
| **LineRAL-380:log10(Dose)** | -0.36718 | 0.1405 | -2.613 | 0.00897 |
| **LineRAL-379:log10(Dose)** | 0.19286 | 0.11074 | 1.742 | 0.08159 |
| **LineRAL-502:log10(Dose)** | 0.23113 | 0.1165 | 1.984 | 0.04725 |
| **LineRAL-75:log10(Dose)** | 0.11279 | 0.13565 | 0.831 | 0.40572 |
| **LineRAL-818:log10(Dose)** | 0.08981 | 0.11412 | 0.787 | 0.43134 |
| **LineRAL-373:log10(Dose)** | 0.38109 | 0.1437 | 2.652 | 0.008 |
| **LineRAL-738:log10(Dose)** | -0.12442 | 0.12613 | -0.986 | 0.32393 |
| **LineRAL-138:log10(Dose)** | -0.34381 | 0.22698 | -1.515 | 0.12984 |
| **LineRAL-765:log10(Dose)** | -0.01657 | 0.10485 | -0.158 | 0.87446 |

**Table S10: Model 5a**

|  | | **Estimate** | **Std. Error** | **t value** | **Pr(>\|t\|)** |
| --- | --- | --- | --- | --- | --- |
| **(Intercept)** | | 4.3825 | 0.111 | 39.469 | <2e-16 |
| **Line.typeS** | | 0.18 | 0.1282 | 1.404 | 0.163 |
| **SexMale** | | 2.505 | 0.1282 | 19.538 | <2e-16 |

**Table S11: Model 5b**

|  | **Estimate** | **Std. Error** | **t value** | **Pr(>\|t\|)** |
| --- | --- | --- | --- | --- |
| **(Intercept)** | 4.65025 | 0.13819 | 33.652 | < 2e-16 |
| **SexMale** | 2.28174 | 0.19542 | 11.676 | < 2e-16 |
| **LineRAL-380** | -1.25285 | 0.19542 | -6.411 | 9.37E-09 |
| **LineRAL-379** | 0.02769 | 0.19542 | 0.142 | 0.88769 |
| **LineRAL-502** | 0.34812 | 0.19542 | 1.781 | 0.07865 |
| **LineRAL-75** | -1.38939 | 0.19542 | -7.11 | 4.37E-10 |
| **LineRAL-818** | 0.42931 | 0.19542 | 2.197 | 0.03093 |
| **LineRAL-373** | -0.15832 | 0.19542 | -0.81 | 0.42026 |
| **LineRAL-738** | -0.56273 | 0.19542 | -2.88 | 0.00511 |
| **LineRAL-138** | 0.42663 | 0.19542 | 2.183 | 0.03196 |
| **LineRAL-765** | 0.3542 | 0.19542 | 1.812 | 0.07367 |
| **SexMale:LineRAL-380** | 0.48808 | 0.27637 | 1.766 | 0.0812 |
| **SexMale:LineRAL-379** | 0.19292 | 0.27637 | 0.698 | 0.48718 |
| **SexMale:LineRAL-502** | 0.34787 | 0.27637 | 1.259 | 0.21179 |
| **SexMale:LineRAL-75** | 0.46169 | 0.27637 | 1.671 | 0.09872 |
| **SexMale:LineRAL-818** | 0.11672 | 0.27637 | 0.422 | 0.67391 |
| **SexMale:LineRAL-373** | 0.34736 | 0.27637 | 1.257 | 0.21247 |
| **SexMale:LineRAL-738** | 0.58875 | 0.27637 | 2.13 | 0.03622 |
| **SexMale:LineRAL-138** | -0.01163 | 0.27637 | -0.042 | 0.96653 |
| **SexMale:LineRAL-765** | -0.29938 | 0.27637 | -1.083 | 0.28195 |

**Table S12: Model 6a**

|  | **Estimate** | **Std. Error** | **t value** | **Pr(>\|t\|)** |
| --- | --- | --- | --- | --- |
| **(Intercept)** | 1.07576 | 0.0805 | 13.363 | < 2e-16 |
| **Line.typeS** | 0.32507 | 0.09295 | 3.497 | 0.000711 |
| **SexMale** | -0.07404 | 0.09295 | -0.796 | 0.427693 |

**Table S13: Model 6b**

|  | **Estimate** | **Std. Error** | **t value** | **Pr(>\|t\|)** |
| --- | --- | --- | --- | --- |
| **(Intercept)** | 1.18529 | 0.20283 | 5.844 | 1.06e-07 |
| **SexMale** | -0.09157 | 0.28685 | -0.319 | 0.75038 |
| **LineRAL-380** | 0.27114 | 0.28685 | 0.945 | 0.34738 |
| **LineRAL-379** | -0.2849 | 0.28685 | -0.993 | 0.3236 |
| **LineRAL-502** | 0.03682 | 0.28685 | 0.128 | 0.89817 |
| **LineRAL-75** | -0.08767 | 0.28685 | -0.306 | 0.76068 |
| **LineRAL-818** | 0.76852 | 0.28685 | 2.679 | 0.00896 |
| **LineRAL-373** | 0.06978 | 0.28685 | 0.243 | 0.80841 |
| **LineRAL-738** | -0.29075 | 0.28685 | -1.014 | 0.31382 |
| **LineRAL-138** | 0.09792 | 0.28685 | 0.341 | 0.73372 |
| **LineRAL-765** | -0.0509 | 0.28685 | -0.177 | 0.8596 |
| **SexMale:LineRAL-380** | -0.23974 | 0.40567 | -0.591 | 0.55619 |
| **SexMale:LineRAL-379** | 0.29471 | 0.40567 | 0.726 | 0.46966 |
| **SexMale:LineRAL-502** | -0.18175 | 0.40567 | -0.448 | 0.65535 |
| **SexMale:LineRAL-75** | -0.07925 | 0.40567 | -0.195 | 0.84561 |
| **SexMale:LineRAL-818** | -0.65997 | 0.40567 | -1.627 | 0.1077 |
| **SexMale:LineRAL-373** | 0.44828 | 0.40567 | 1.105 | 0.27245 |
| **SexMale:LineRAL-738** | 0.21158 | 0.40567 | 0.522 | 0.60341 |
| **SexMale:LineRAL-138** | 0.44908 | 0.40567 | 1.107 | 0.2716 |
| **SexMale:LineRAL-765** | -0.06761 | 0.40567 | -0.167 | 0.86806 |

**Table S14: Model 7a**

|  | **Estimate** | **Std. Error** | **t value** | **Pr(>\|t\|)** |
| --- | --- | --- | --- | --- |
| **(Intercept)** | 13.3054 | 0.1992 | 66.788 | < 2e-16 |
| **Line.typeS** | -0.8116 | 0.23 | -3.528 | 0.000641 |
| **SexMale** | -1.6125 | 0.23 | -7.01 | 3.19E-10 |

**Table S15: Model 7b**

|  | **Estimate** | **Std. Error** | **t value** | **Pr(>\|t\|)** |
| --- | --- | --- | --- | --- |
| **(Intercept)** | 13.9901 | 0.4648 | 30.097 | < 2e-16 |
| **SexMale** | -3.5482 | 0.6574 | -5.397 | 6.72E-07 |
| **LineRAL-380** | -2.3399 | 0.6574 | -3.559 | 0.000629 |
| **LineRAL-379** | -0.15 | 0.6574 | -0.228 | 0.820059 |
| **LineRAL-502** | -0.2113 | 0.6574 | -0.321 | 0.748748 |
| **LineRAL-75** | -0.8833 | 0.6574 | -1.344 | 0.182869 |
| **LineRAL-818** | -0.6402 | 0.6574 | -0.974 | 0.333075 |
| **LineRAL-373** | -1.5613 | 0.6574 | -2.375 | 0.019939 |
| **LineRAL-738** | -1.4879 | 0.6574 | -2.263 | 0.02632 |
| **LineRAL-138** | -1.4337 | 0.6574 | -2.181 | 0.032132 |
| **LineRAL-765** | -2.1971 | 0.6574 | -3.342 | 0.001266 |
| **SexMale:LineRAL-380** | 2.8309 | 0.9297 | 3.045 | 0.003149 |
| **SexMale:LineRAL-379** | 0.9933 | 0.9297 | 1.068 | 0.288516 |
| **SexMale:LineRAL-502** | 1.6486 | 0.9297 | 1.773 | 0.079985 |
| **SexMale:LineRAL-75** | 3.3668 | 0.9297 | 3.621 | 0.000512 |
| **SexMale:LineRAL-818** | 2.0374 | 0.9297 | 2.192 | 0.031324 |
| **SexMale:LineRAL-373** | 1.7459 | 0.9297 | 1.878 | 0.064035 |
| **SexMale:LineRAL-738** | 2.2882 | 0.9297 | 2.461 | 0.015998 |
| **SexMale:LineRAL-138** | 1.4566 | 0.9297 | 1.567 | 0.121113 |
| **SexMale:LineRAL-765** | 2.9894 | 0.9297 | 3.216 | 0.001879 |

**Table S16: Model 8a**

|  | **Estimate** | **Std.Error** | **tvalue** | **Pr(>\|t\|)** |
| --- | --- | --- | --- | --- |
| **(Intercept)** | 18.172 | 4.629 | 3.926 | 0.000163 |
| **Line.typeS** | -6.191 | 6.546 | -0.946 | 0.34663 |
| **SexMale** | -14.815 | 6.546 | -2.263 | 0.025875 |
| **Line.typeS:SexMale** | 16.327 | 9.257 | 1.764 | 0.080973 |

**Table S17: Model 8b**

|  | **Estimate** | **Std. Error** | **t value** | **Pr(>\|t\|)** |
| --- | --- | --- | --- | --- |
| **(Intercept)** | 5.3178 | 8.8864 | 0.598 | 0.55125 |
| **SexMale** | -3.6795 | 12.5673 | -0.293 | 0.77044 |
| **LineRAL-380** | 0.4886 | 12.5673 | 0.039 | 0.96908 |
| **LineRAL-379** | 52.2282 | 12.5673 | 4.156 | 8.07E-05 |
| **LineRAL-502** | -3.2022 | 12.5673 | -0.255 | 0.79953 |
| **LineRAL-75** | 17.4582 | 12.5673 | 1.389 | 0.16864 |
| **LineRAL-818** | 6.0237 | 12.5673 | 0.479 | 0.63302 |
| **LineRAL-373** | 1.4227 | 12.5673 | 0.113 | 0.91015 |
| **LineRAL-738** | -2.212 | 12.5673 | -0.176 | 0.86073 |
| **LineRAL-138** | 29.0024 | 12.5673 | 2.308 | 0.0236 |
| **LineRAL-765** | -3.6204 | 12.5673 | -0.288 | 0.77403 |
| **SexMale:LineRAL-380** | 13.2762 | 17.7729 | 0.747 | 0.45726 |
| **SexMale:LineRAL-379** | -48.1163 | 17.7729 | -2.707 | 0.00829 |
| **SexMale:LineRAL-502** | 2.8207 | 17.7729 | 0.159 | 0.8743 |
| **SexMale:LineRAL-75** | -12.1154 | 17.7729 | -0.682 | 0.49741 |
| **SexMale:LineRAL-818** | 32.59 | 17.7729 | 1.834 | 0.07042 |
| **SexMale:LineRAL-373** | 6.5956 | 17.7729 | 0.371 | 0.71154 |
| **SexMale:LineRAL-738** | 1.7353 | 17.7729 | 0.098 | 0.92247 |
| **SexMale:LineRAL-138** | -29.5377 | 17.7729 | -1.662 | 0.10044 |
| **SexMale:LineRAL-765** | 3.033 | 17.7729 | 0.171 | 0.86493 |
