## Supplementary Figure for "The restriction factor *pastrel* is associated with host vigor, viral titer, and variation in disease tolerance during Drosophila C Virus infection"

### Supplementary figures

**Figures S1 –S17**

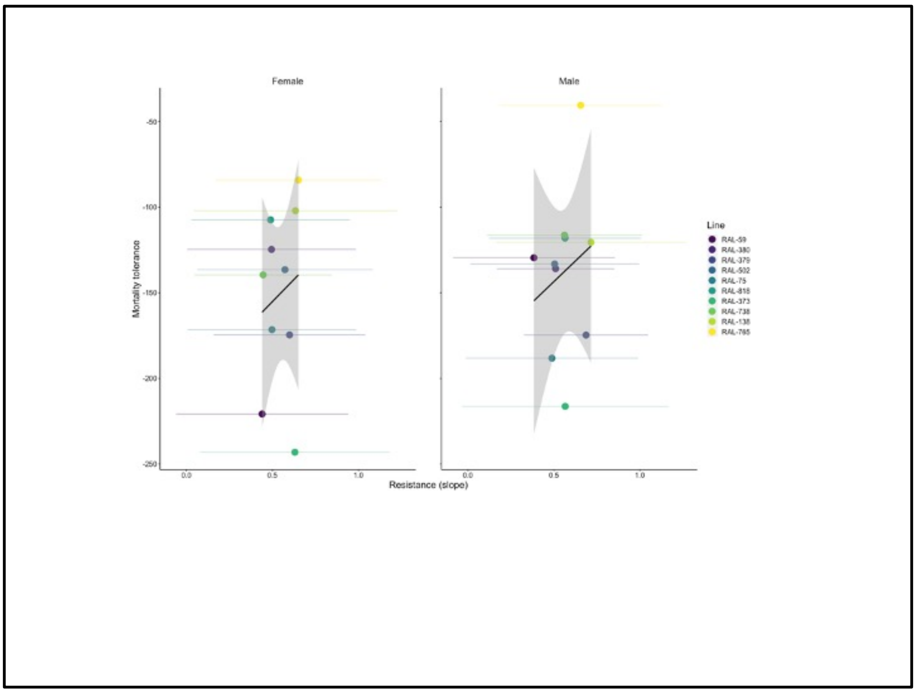


**Figure S1- Correlation between mortality tolerance and resistance**

**Females**
Kendall's rank correlation tau
T = 27, p-value = 0.4843

**Males**
Kendall's rank correlation tau
$Slope_titre and tls_m$Nl_life
T = 25, p-value = 0.7275

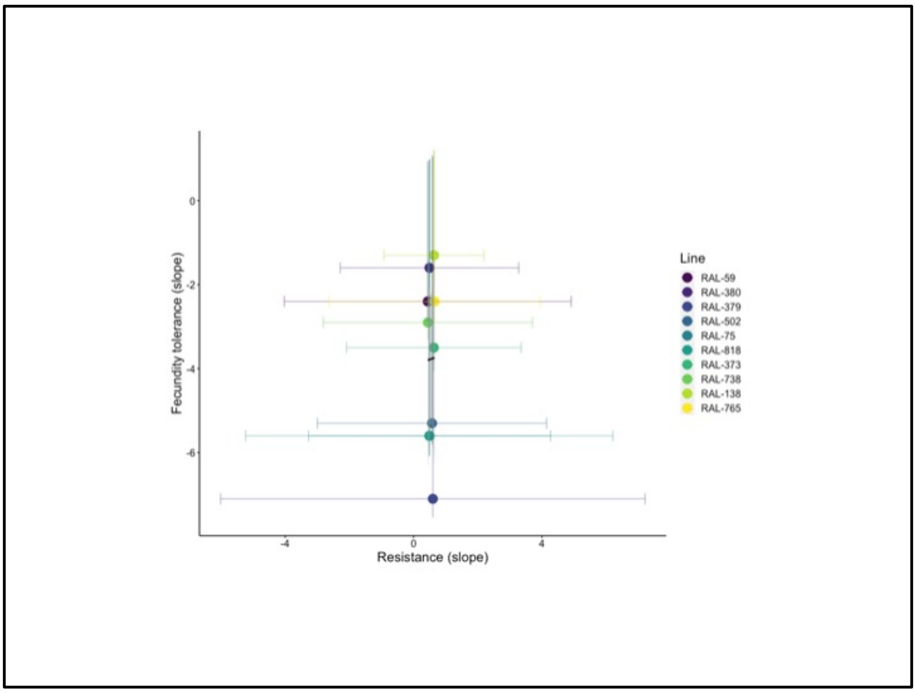


**Figure S2 - Correlation between fecundity tolerance and resistance**

Kendall's rank correlation tau
z = 0.2705, p-value = 0.7868

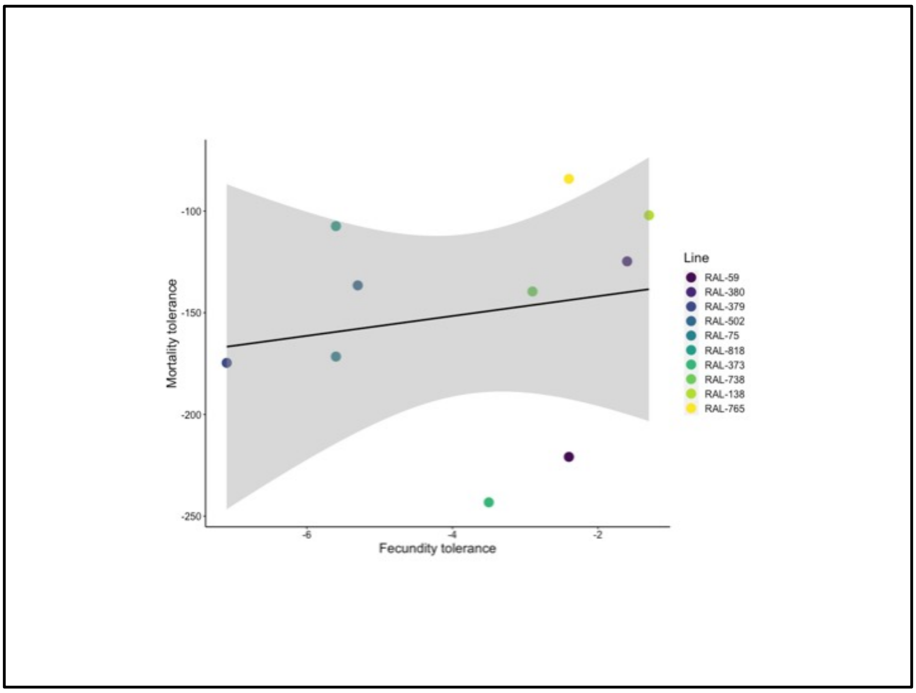


**Figure S3 - no trade-off between mortality tolerance and fecundity tolerance**
Kendall's rank correlation tau

z = 1.1722, p-value = 0.2411
tau = 0.2955309

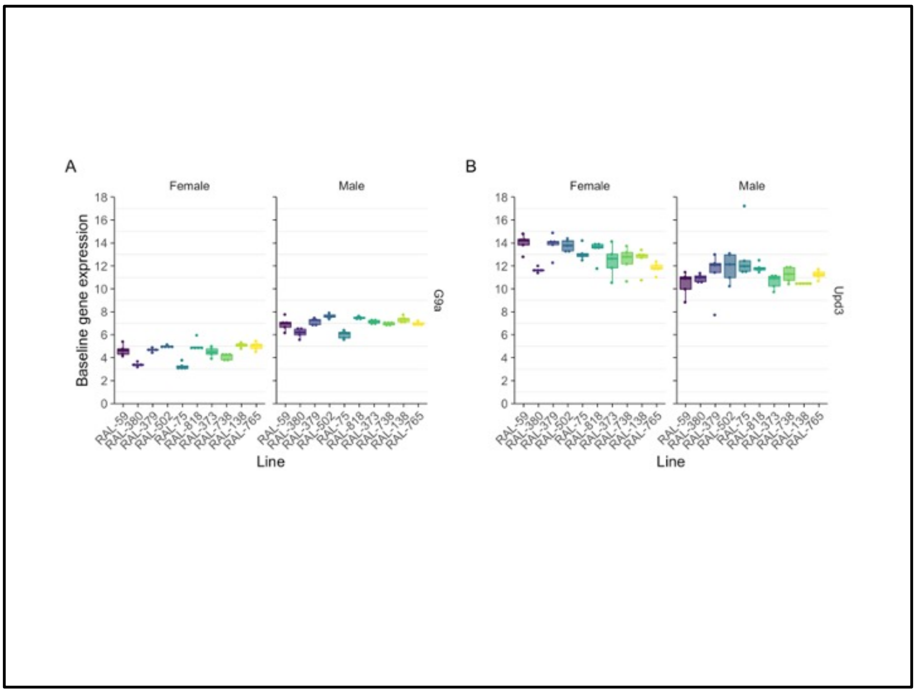


**Figure S4 Baseline gene expression (pre-infection)relative to rp49. For statistics see Table 4.**

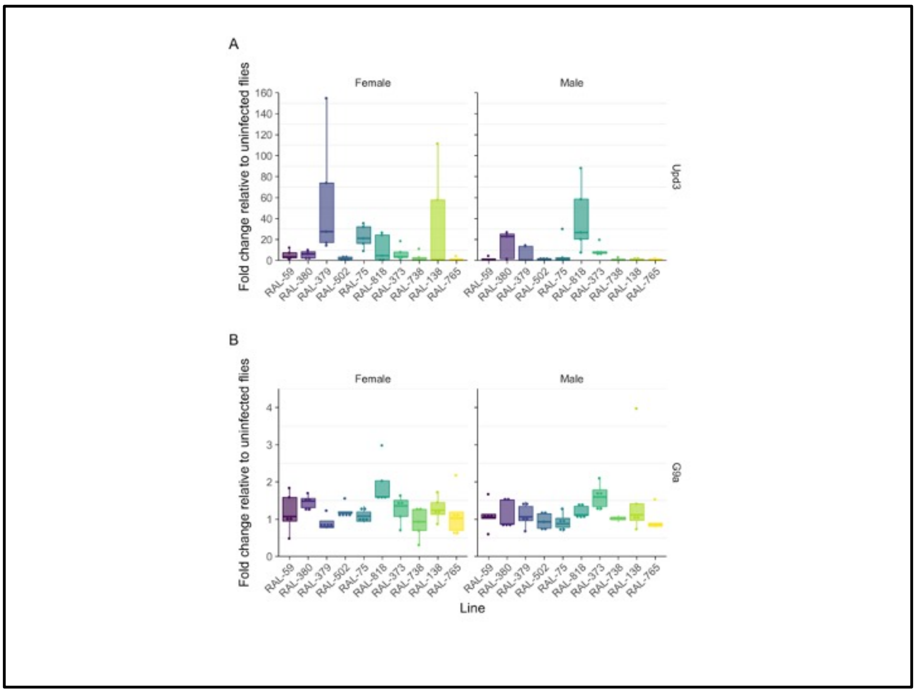


**Figure S5 Gene expression in DCV infected flies relative to uninfected flies.** For statistics see Table 4.

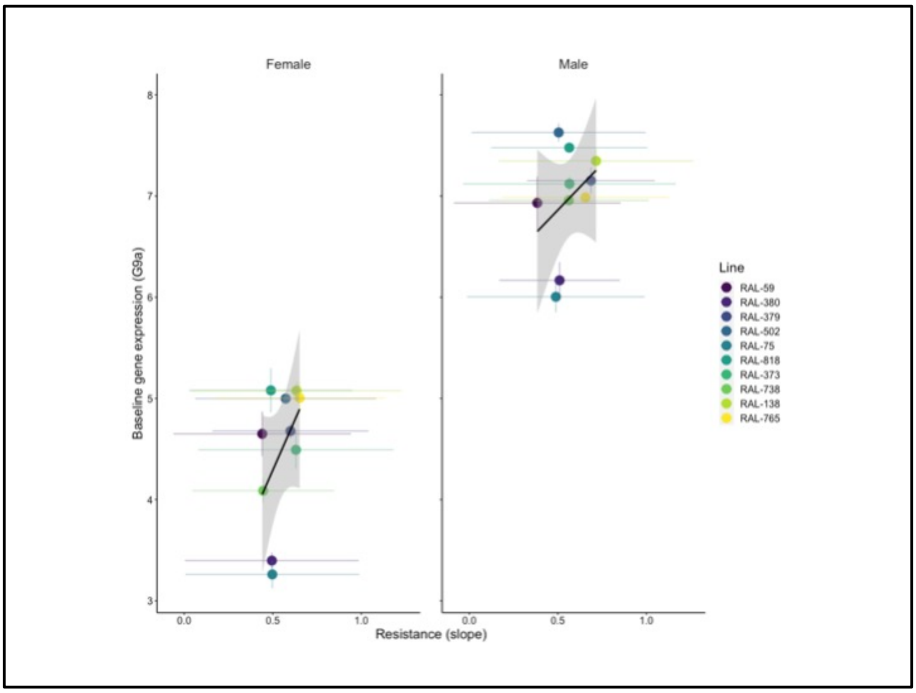


**Figure S6 – Correlation between baseline expression of G9a and resistance.**

**Females**
Kendall's rank correlation tau
T = 27, p-value = 0.4843

**Males**
Kendall's rank correlation tau
T = 31, p-value = 0.1557

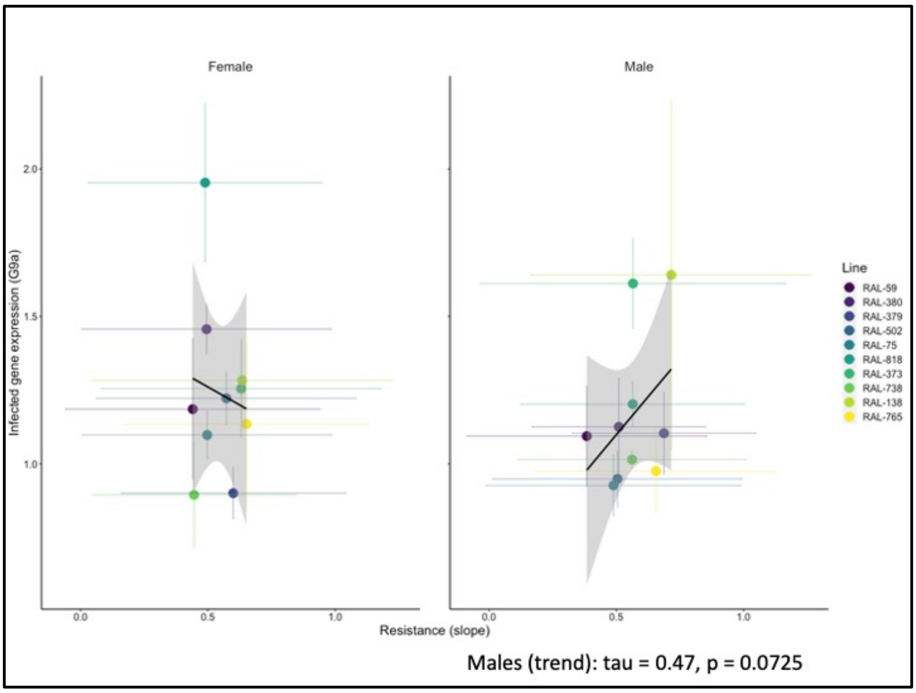


**Figure S7- Correlation between infected expression of G9a and resistance**

**Females**
Kendall's rank correlation tau
T = 23, p-value = 1

**Males**
Kendall's rank correlation tau
T = 33, p-value = 0.07255
tau
0.4666667

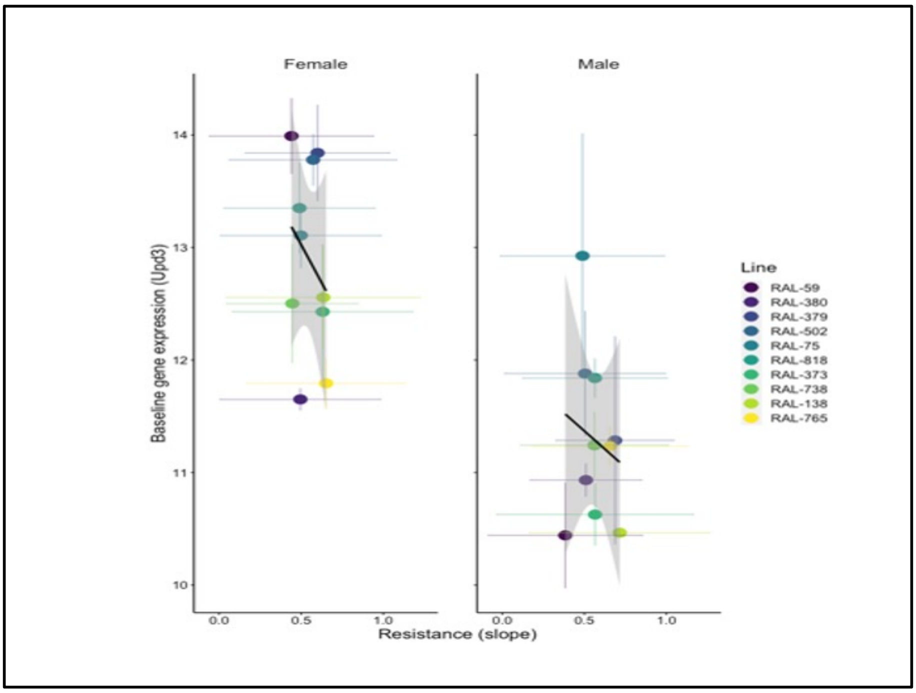


**FigureS8- Correlation between baseline expression of Upd3 and resistance**

**Females**
Kendall's rank correlation tau
T = 17, p-value = 0.3807

**Males**
Kendall's rank correlation tau
T = 18, p-value = 0.4843

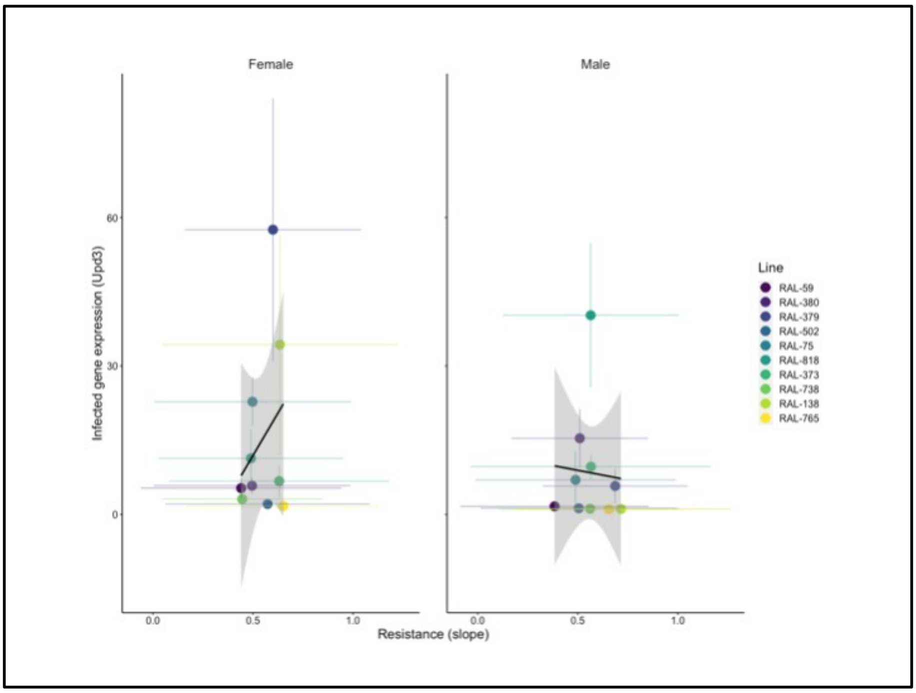


**Figure S9 - Correlation between infected expression of Upd3 and resistance**

**Females**
Kendall's rank correlation tau
T = 25, p-value = 0.7275

**Males**
Kendall's rank correlation tau
T = 18, p-value = 0.4843

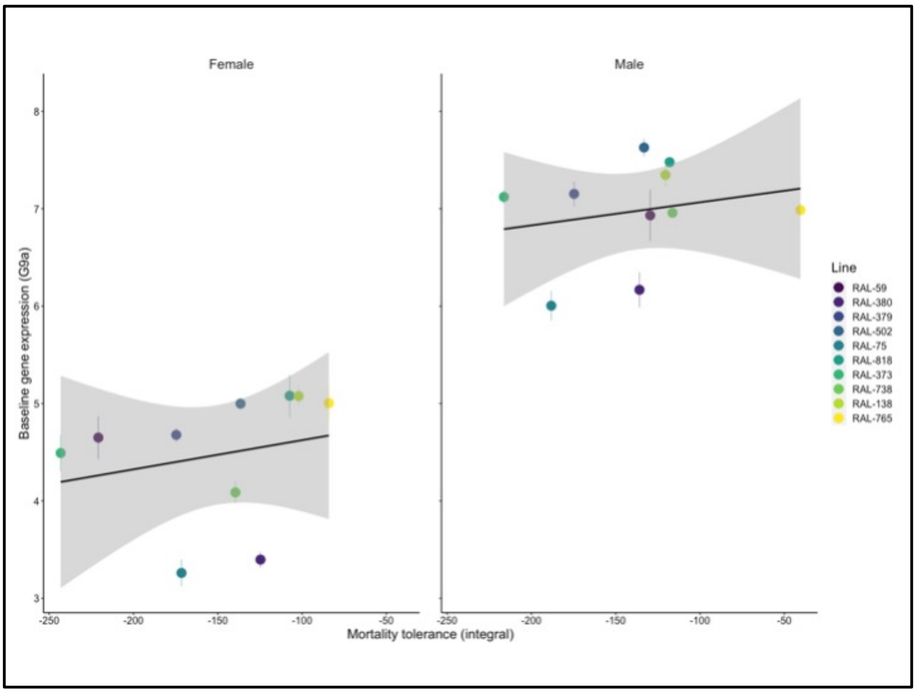


**Figure S10- Correlation between baseline expression of G9a and mortality tolerance**

**Females**
Kendall's rank correlation tau
T = 27, p-value = 0.4843

**Males**
Kendall's rank correlation tau
T = 31, p-value = 0.1557

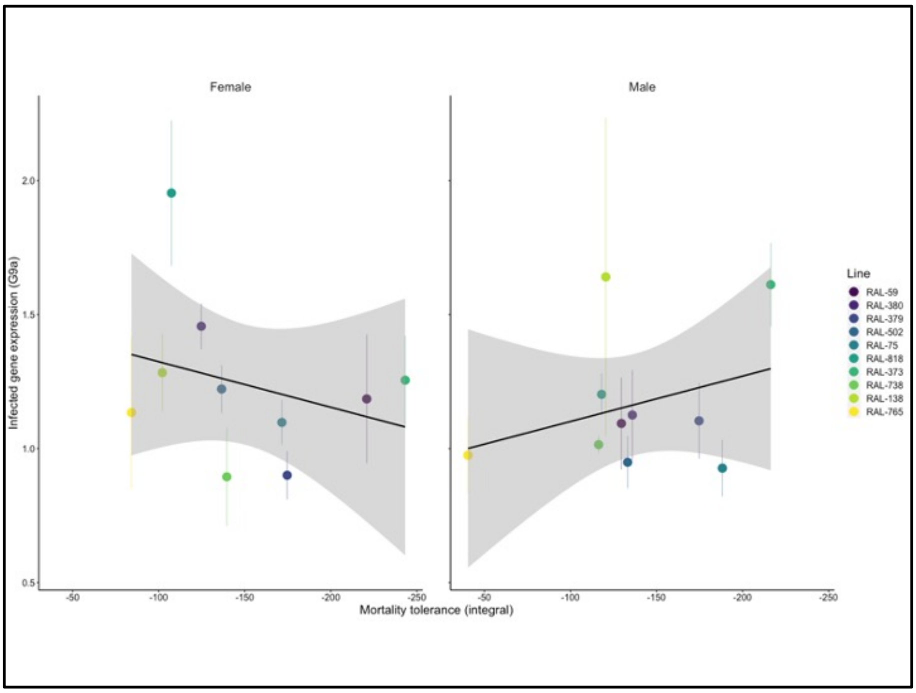


**Figure S11 - Correlation between infected expression of G9a and mortality tolerance**

**Females**

Kendall's rank correlation tau

T = 27, p-value = 0.4843

**Males**

Kendall's rank correlation tau

T = 21, p-value = 0.8618

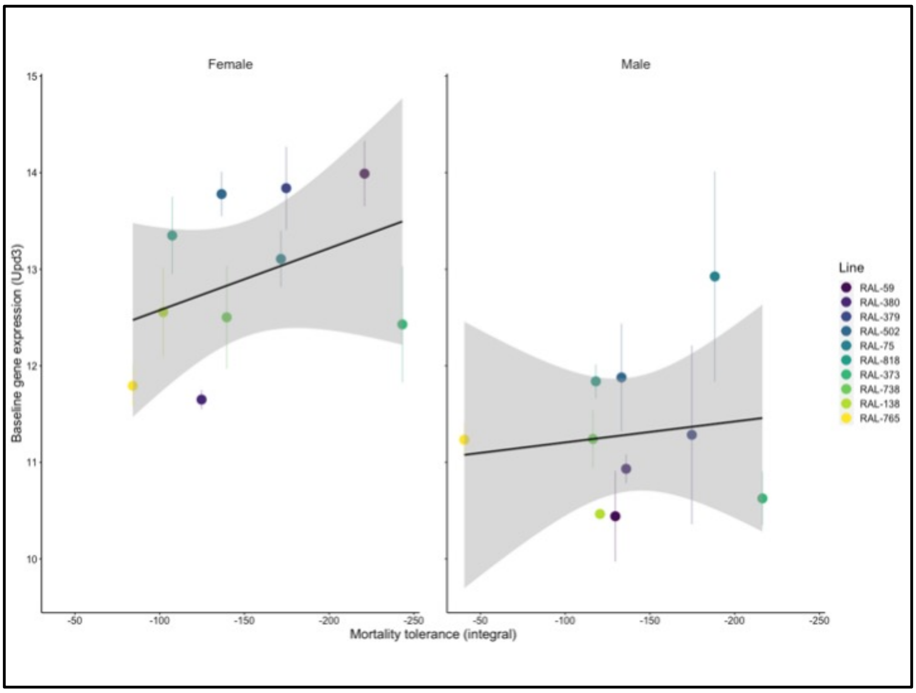


**Figure S12 - Correlation between baseline expression of Upd3 and mortality tolerance**

**Females**

Kendall's rank correlation tau

T = 15, p-value = 0.2164

**Males**

Kendall's rank correlation tau

T = 20, p-value = 0.7275

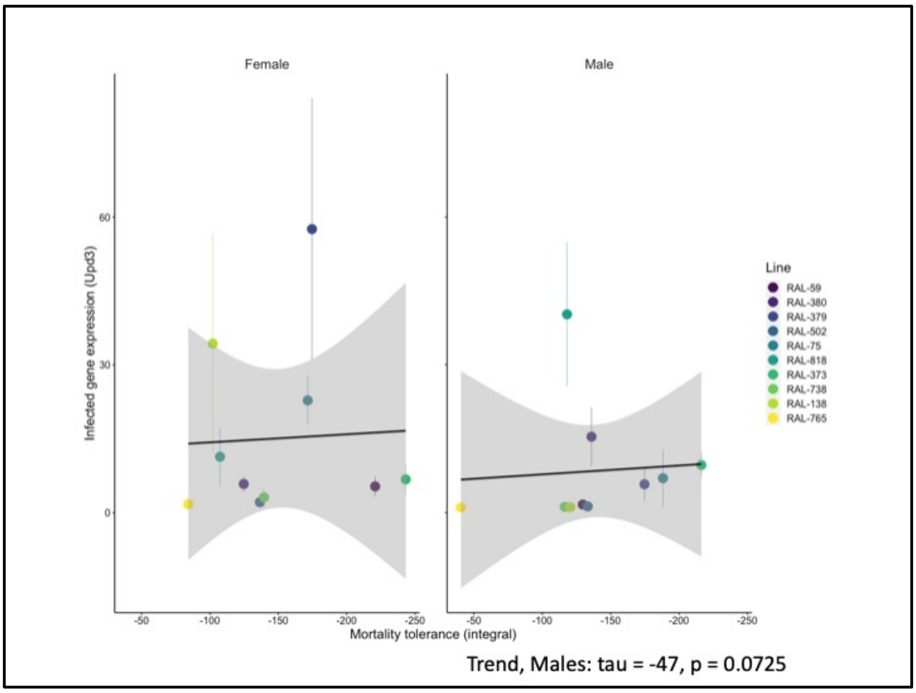


**Figure S13 - Correlation between infected expression of Upd3 and mortality tolerance**

**Females**

Kendall's rank correlation tau

T = 19, p-value = 0.6007

**Males**

Kendall's rank correlation tau

T = 12, p-value = 0.07255

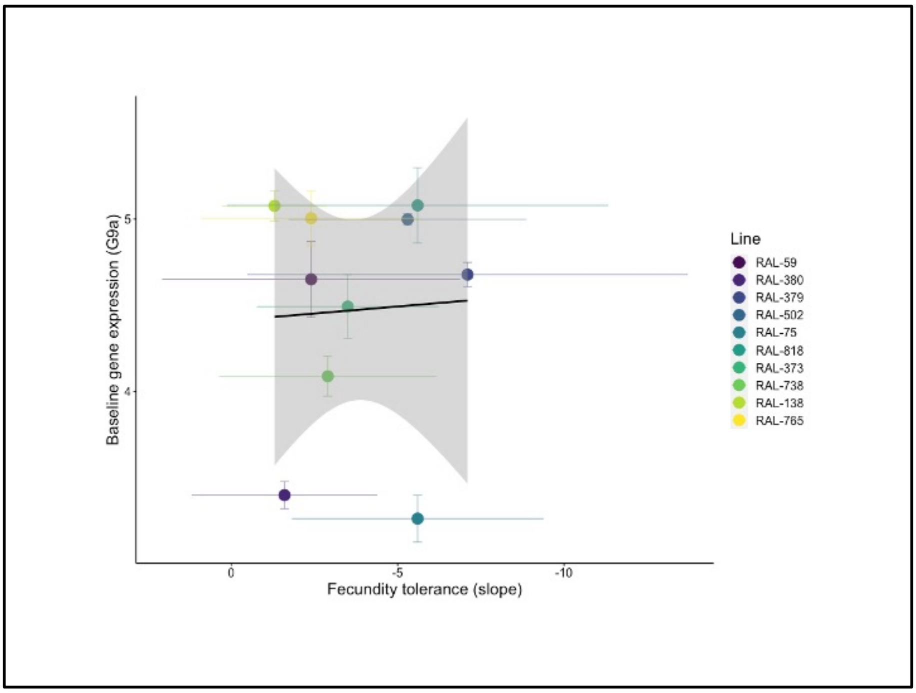


**Figure S14 - Correlation between baseline expression of G9a and fecundity tolerance**

z = 0.090167, p-value = 0.9282

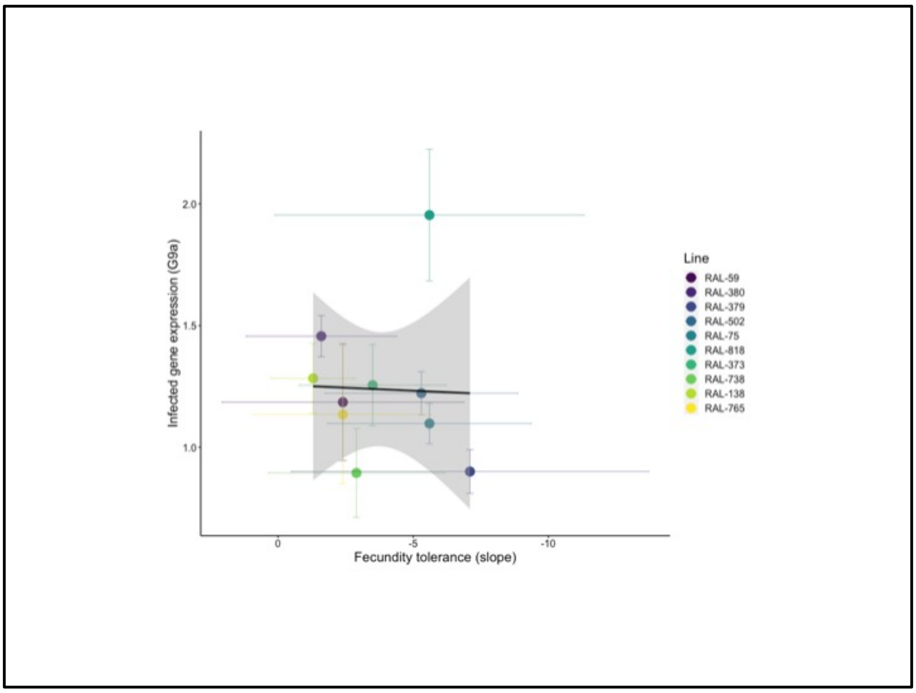


**Figure S15 - Correlation between infected expression of G9a and fecundity tolerance**

Kendall's rank correlation tau
z = 0.99184, p-value = 0.3213

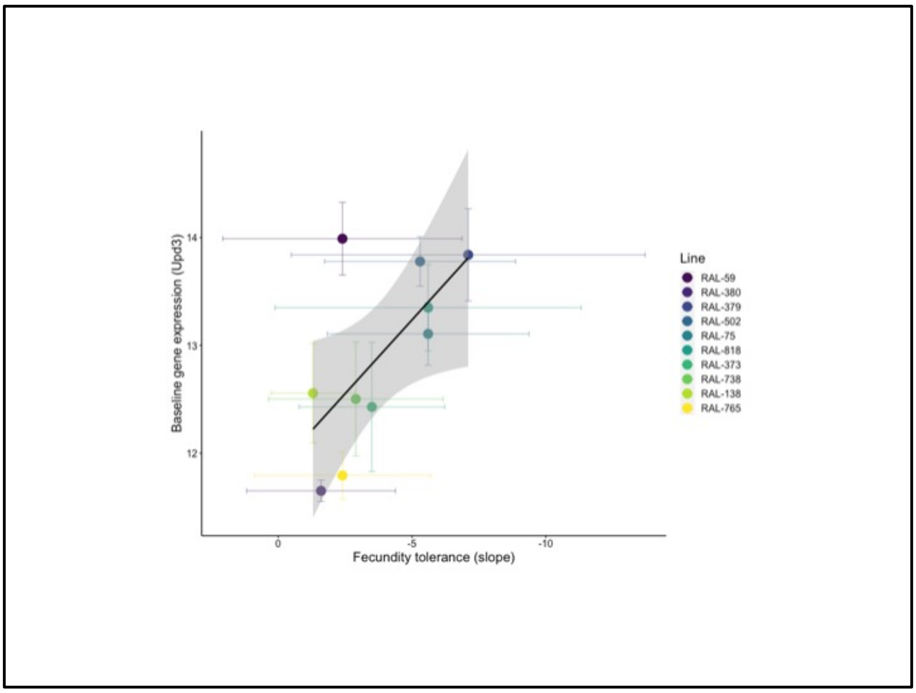


**Figure S16 - Correlation between baseline expression of Upd3 and fecundity tolerance**

Kendall's rank correlation tau
z = -1.5328, p-value = 0.1253

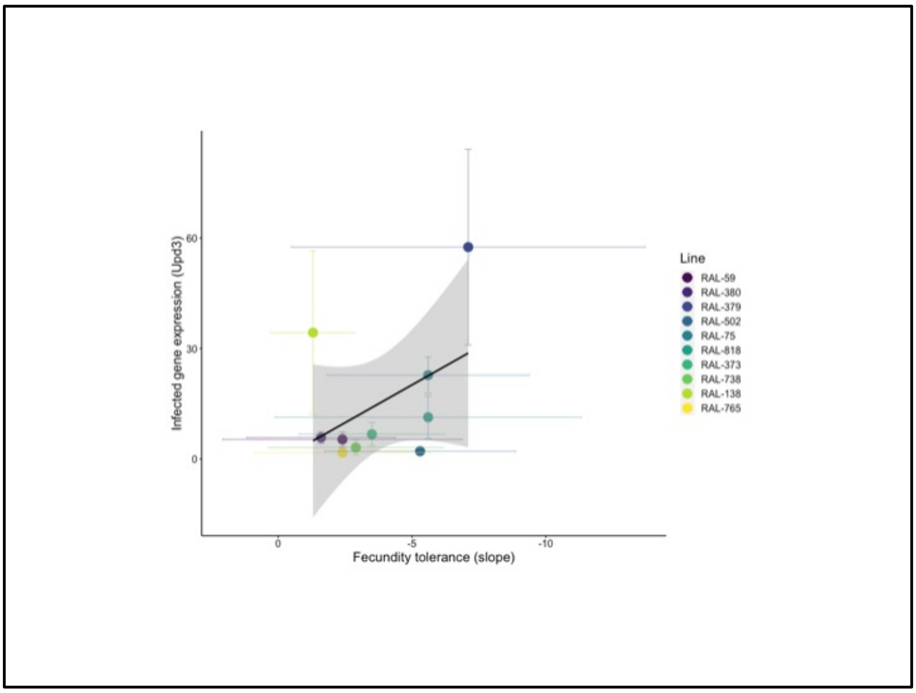


**Figure S17** - **Correlation between infected expression of Upd3 and fecundity tolerance**

Kendall's rank correlation tau
z = -0.99184, p-value = 0.3213
